# Transcriptional activation forms insulation boundaries through cohesin-independent decompaction and cohesin-dependent barrier formation

**DOI:** 10.64898/2026.11.24.768880

**Authors:** Jamie A. Drayton, Asmita Jha, Jack Toppen, Sean Manley, Anders S. Hansen

## Abstract

How 3D genome structure regulates transcription has been intensely studied, but how transcription regulates 3D genome structure remains poorly understood. To elucidate this relationship, we integrate Region Capture Micro-C and simultaneous live-cell imaging of CTCF chromatin looping and inducible nascent transcription at the *Fbn2* locus with 3D polymer modeling and perturbation experiments. We find that transcription activation induces cohesin-independent insulation boundary formation at the *Fbn2* promoter, locus decompaction, and faster chromatin dynamics. We also show that transcriptional bursts interfere with CTCF loop formation in living cells. Using 3D polymer modeling to distinguish plausible mechanistic models, we find that transcription-induced gene body decompaction can explain increased mobility and cohesin-independent insulation, while a polymerase-associated barrier to loop extrusion can explain transcription-linked looping changes. Together, these results reveal cohesin-dependent and -independent mechanisms through which transcription can establish insulation and regulate 3D genome structure.

## INTRODUCTION

The mammalian genome is organized into complex nested structures across a range of length scales^1^, from chromosome territories^2,3^ and A/B-compartments^4^, through topologically-associating domains (TADs)^5–7^ and loops^8–10^ at intermediate scales (∼50kb-1Mb), and down to individual nucleosomes. TADs are formed by cohesin, which loads onto chromatin and dynamically extrudes it into loops typically bounded by CTCF^11–20^. Beyond CTCF, TAD boundaries are also enriched in active promoters^6^. Intermediate-scale 3D organization is key to gene regulation, for example by modulating contacts between gene promoters and the enhancers that regulate their expression^17,21–27^.

However, it is less clear how transcription regulates genome organization at the level of TADs and loops. Many possible models of this relationship have been proposed; some depend on interaction between transcription and the cohesin protein complex, while others are independent of cohesin. Some studies indicate that transcribing RNA polymerase II (RNAPII) may interfere with loop extrusion by blocking and relocating cohesins^28–36^, potentially erasing TAD boundaries entirely^31^; others suggest regulatory elements such as promoters or enhancers may serve as targeted loading sites for cohesin^37–45^. Cohesin-independent models of transcription-associated 3D genome changes primarily focus on modulation of chromatin compaction in and around active genes. However, the direction of this modulation is debated: activation of transcription has been linked to both compaction^46–48^ and decompaction^31,48–50^ of active loci in different works. Thus, despite many studies, it remains unclear whether transcription reinforces or antagonizes intermediate-scale 3D genome organization.

The discrepancies in the literature are partially due to technical limitations. Direct super-resolved simultaneous visualization of CTCF/cohesin-mediated looping and inducible and programmable nascent transcription has not yet been achieved. This is important because transcription is not constitutive, but occurs in infrequent bursts^51,52^, such that any bulk method averages over both transcribing and non-transcribing states. Furthermore, while live-cell imaging reports absolute distances, genomics approaches are inherently normalized and relative. For example, matrix balancing of 3C-based data makes it challenging to distinguish between compaction and decompaction^49^. Here, we overcome these limitations by integrating multiple methods using the same model system of a synthetically inducible gene and its surrounding TAD. We combine super-resolution live-cell imaging (SRLCI)^15–17,53^ to directly probe the impact of transcriptional machinery on chromatin dynamics during inducible nascent transcription with high-resolution Region-Capture Micro-C (RCMC)^54^ to profile locus-wide 3D genome changes upon transcriptional activation and perturbations to CTCF and cohesin. We then conduct 3D polymer simulations to assess which mechanistic models best explain our comprehensive experimental dataset. Our results reveal that promoters and gene bodies can form 3D insulation boundaries in both cohesin-dependent and cohesin-independent manners. First, gene body decompaction forms cohesin-independent insulation boundaries. Second, RNAPII mediates cohesin-dependent 3D insulation by serving as a moving barrier to loop extrusion. Taken together, our results provide a unified model bridging cohesin-dependent and -independent mechanisms of transcription-mediated 3D genome insulation.

## RESULTS

### A system for probing looping-transcription interplay with high spatiotemporal resolution

To study the interactions between cohesin-mediated looping and transcriptional machinery, we elected to study the *Fbn2* gene and its surrounding locus in mESCs as our model locus. The *Fbn2* TAD, which is simple and well-characterized, is a CTCF-dependent ∼505kb TAD^55^ anchored by 2 CTCF sites on the left (L1 and L2, separated by 65 kb) and a single CTCF site on the right (R1). It contains a single ∼201.5kb gene, *Fbn2*, that is transcriptionally inactive in mESCs. All prior 3D genomics studies in mESCs that we examined – including Hi-C, Micro-C, and RCMC performed in different mESC lineages grown under different conditions and studied by different groups – consistently find that the *Fbn2* region shows the key features of a TAD or loop domain: a triangle along the diagonal anchored by one major loop (L1-R1) and one minor loop (L2-R1) (Fig. 1a)^9,11,15,54,56–58^ and that the *Fbn2* TAD is lost upon CTCF or cohesin depletion^15,54,56,57^. Furthermore, prior studies have shown that the *Fbn2* TAD is fully looped (CTCF sites connected by one or more cohesins) ∼5% of the time^15,58^, making it one of the strongest CTCF loops in the mESC genome^58^, and in a partially extruded configuration (cohesin loaded somewhere within the TAD) ∼92% of the time^15^.

**Figure 1:**
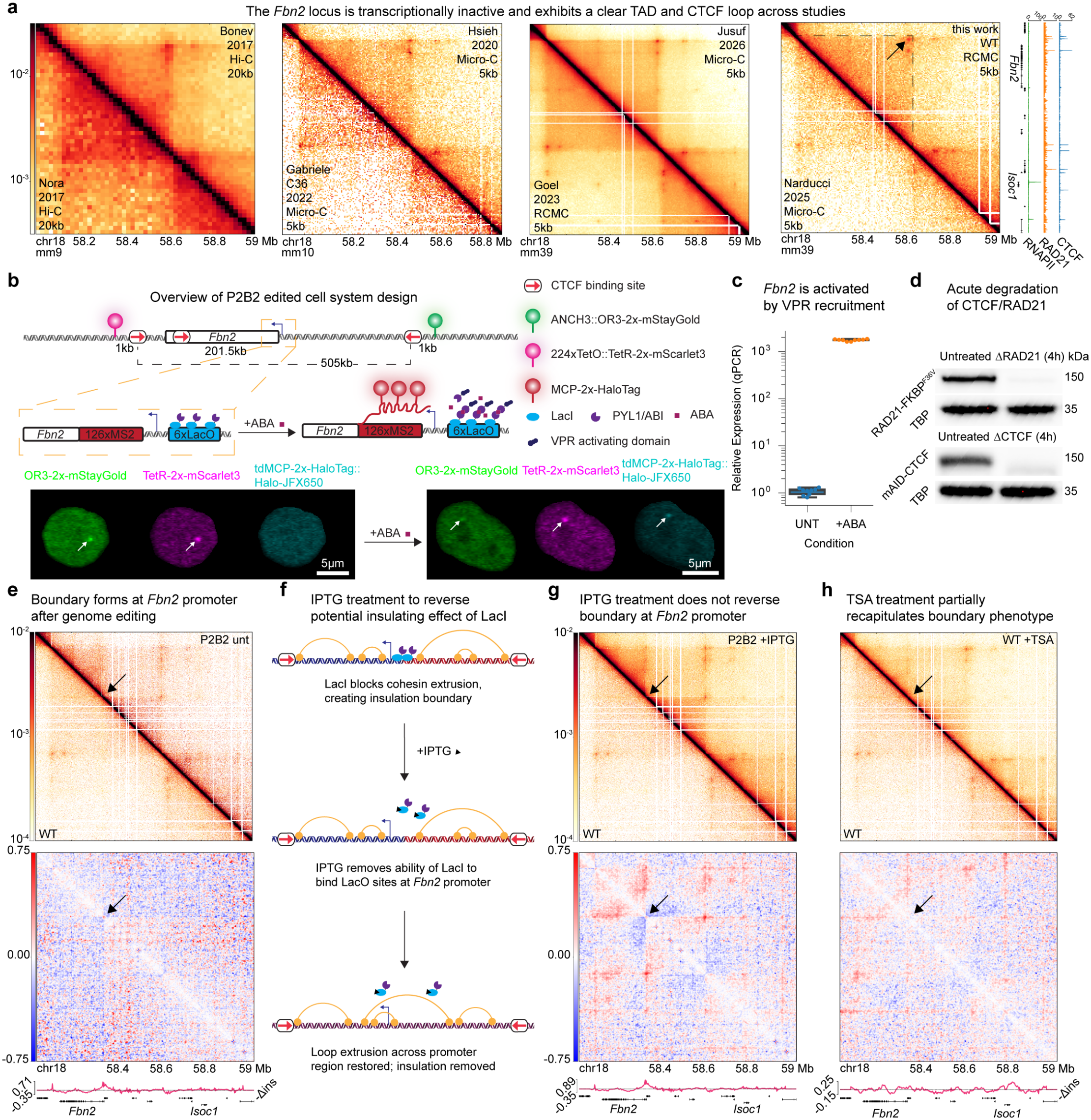
A system for characterizing spatiotemporal effects of transcriptional activation on genome organization. **a**, Contact map visualizations of the *Fbn2* locus from several prior studies^9,11,15,54,56–58^ and our own work. The *Fbn2* TAD is indicated with dashed lines, and the CTCF-CTCF loop with an arrow. Genomics method (Hi-C, Micro-C, or RCMC) and displayed bin size are indicated on each map. Right: ChIP-seq signals for CTCF (GSE90994), RAD21 (GSE90994) and RNAPII (GSE58019) across the *Fbn2* locus. **b**, Schematic representation of genome-edited cell system and example single z-slices from live-cell images of edited cells. Dots representing DNA loci and an active MS2 burst are indicated with arrows. **c**, qPCR expression of *Fbn2* and the antisense transcript from the *Fbn2* promoter. Values are normalized to mean expression in untreated P2B2 cells. **d**, Western blots showing depletion of RAD21 and CTCF after acute (4 hours) degron treatment. **e**, Top: Contact map visualization of RCMC data from P2B2 cells (top right) and WT JM8.N4 cells (bottom left). Middle: Map showing P2B2/WT contact ratio. Bottom: Track of differential insulation score (WT - P2B2). **f**, Schematic overview of IPTG treatment. **g**, As **e**, for IPTG-treated (7 days) P2B2 cells. **h**, As **e**, for TSA-treated (4 hours) WT cells.

Thus, we reasoned that the *Fbn2* TAD would be ideally suited for a bottom-up synthetic biology approach to study the impact of transcription on 3D genome organization. Rather than re-using our previously described cell lines for live-cell imaging of the *Fbn2* TAD reported in Gabriele *et al*.^15^, which already contain Halo- and SNAP_f_-tagged CTCF and RAD21, we re-engineered wild-type JM8.N4 mESCs^59^ using the same genome-editing constructs. We began by inserting a 224xTetO array ∼5kb upstream of the L1 CTCF site and an ANCH3 array^60^ ∼5kb downstream of the R1 CTCF site. To improve localization precision in live-cell imaging^61^, we then used Cas9-mediated genome editing to shorten the tether distances between the 224xTetO and ANCH3 arrays at either end of the TAD and their respective CTCF sites from ∼5kb to ∼1kb on each side (Fig. S1a)^15,61,62^. To enable inducible activation of *Fbn2* transcription, we inserted a short 6xLacO array ∼500bp upstream of the promoter of *Fbn2*; we then inserted an activation construct consisting of LacI-ABI and PYL1-VPR fusion proteins^63,64^ at the *Hipp11* safe harbor locus^65,66^ (Fig. 1b, S1a). In the presence of abscisic acid (ABA), ABI and PYL1 form a complex with each other, bringing the synthetic activator VPR to the *Fbn2* promoter and robustly activating transcription by >1,000-fold (Fig. 1b,c, S1b)^63^. Next, we inserted a 126xMS2 array^17,67^ into the first exon of *Fbn2*, ∼400bp downstream of the TSS (Fig. 1b), to visualize nascent transcripts at the locus. To investigate the contributions of loop extrusion machinery to activation-associated 3D genome changes, we tagged *Ctcf* and cohesin subunit *Rad21* with orthogonal degron tags (mAID^68^ and FKBP12^F36V^[dTag]^69^, respectively) (Fig. S1a). Acute depletion allowed us to minimize secondary effects and calibrate the Bayesian Inference of Looping Dynamics (BILD) loop quantification algorithm previously described by our group^15^. We confirmed that both proteins could be acutely degraded after 4 hours of treatment with the respective small molecules (Fig. 1d). Finally, we co-expressed TetR-2x-mScarlet3^70^, OR3-2x-StayGold^60,71^, and tdMCP-2x-HaloTag^67,72,73^ constructs to enable simultaneous live-cell imaging of loop extrusion between the TAD anchors and active transcription at *Fbn2*, and confirmed that dots corresponding to the TAD anchors and active bursts were visible by microscopy (Fig. 1b, S1a). The final edited cell line (hereafter clone P2B2) allowed us to simultaneously visualize TAD anchor dynamics and nascent transcription and to control both transcription of *Fbn2* and perturbations of loop extrusion dynamics.

### Unexpected insulation at the *Fbn2* promoter after genome editing

Unexpectedly, RCMC^54^ of the edited *Fbn2* locus in P2B2 cells revealed a “boundary” phenotype characterized by increased insulation at the *Fbn2* promoter (i.e., reduced frequency of contacts between regions on opposite sides of the promoter), even when cells were not treated with ABA and *Fbn2* was not being actively transcribed (Fig. 1e). This feature has not been observed in any prior 3D genomics studies of this locus that we examined, including Micro-C performed on the C36 genome-edited line used in Gabriele *et al*. to originally visualize *Fbn2* TAD dynamics^15^, or our own Micro-C performed on wild-type mESCs (Fig. 1a)^9,11,15,54,56–58^. We sought to investigate potential mechanisms underlying this effect. We first hypothesized that LacI-ABI binding to the 6xLacO operators inserted near the *Fbn2* promoter might block cohesin-mediated loop extrusion and thus form an insulation barrier. To test this, we treated P2B2 cells with IPTG for 7 days to remove binding of LacI-ABI and performed RCMC. This treatment did not remove the promoter insulation boundary (Fig. 1f,g), consistent with prior work that has also found no evidence of LacI blocking cohesin extrusion^16^. As boundaries like this have recently been reported to form after Cas9-mediated genome editing in other systems^74,75^, we also considered whether genome editing could have affected chromatin accessibility at the *Fbn2* promoter. Indeed, ATAC-seq showed an increase in accessibility at the *Fbn2* promoter in P2B2 cells compared to wild-type (Fig. S2a). To correct for possible biases in RCMC induced by the increase in accessibility, we re-analyzed our contact maps using neighbor balancing^49^, which corrects for non-uniform contact frequencies along the linear genome, such as might be induced by a dramatic accessibility transition. The insulation boundary persisted in maps made from neighbor-balanced data, ruling out technical bias due to accessibility as an explanation (Fig. S2b).

Motivated by the increased accessibility of the *Fbn2* promoter in the P2B2 clone, we next sought to examine if epigenomic state changes might be sufficient to partially reproduce the insulation boundary. To this end, we treated unedited wild-type mESCs with trichostatin A (TSA, a global HDAC inhibitor) to induce hyperacetylation, mimicking more active environments, or A-485 (a p300 inhibitor) to induce hypoacetylation, mimicking more repressed environments (Fig. S2c)^76,77^. TSA-treated cells partially reproduced the insulation boundary coupled with increased H3K27Ac levels at the *Fbn2* promoter (Fig. 1h). A-485 treatment had minimal effect on the normally inactive *Fbn2* locus (Fig. S2d,e). Though the insulation boundary discussed here appears linked to some other phenomenon not directly connected to the active/inactive state of the promoter, as *Fbn2* is not expressed without ABA treatment, these results suggest that epigenomic changes of inactive regulatory regions can trigger 3D genome re-organization. Thus, we aimed to characterize the 3D genome effects of inducing activation at the *Fbn2* locus.

### Transcriptional activation strengthens insulation independently of loop extrusion

To probe the effects of active transcription on the 3D genome, we treated P2B2 cells with ABA overnight to activate transcription at *Fbn2* and performed RCMC (Table S1, Fig. S3a). Cells with active transcription showed stronger insulation across the *Fbn2* promoter (Fig. 2a). Remarkably, this effect was consistent in strength even in cells where CTCF, RAD21, or both had been degraded (Fig. 2a). This indicates that the increase in *Fbn2* promoter insulation upon transcription induction is due to a mechanism unrelated to cohesin-mediated loop extrusion and TAD formation, which require cohesin and CTCF. Although targeted cohesin-loading at promoters has been proposed^37–40^, we observed no signs of targeted cohesin loading at the active *Fbn2* promoter, which would have been expected to result in a strong “jet” or “fountain” perpendicular to the diagonal in contact maps^41–43,45,78,79^. As active regulatory elements have been previously reported to be able to form punctate loop interactions, we sought to characterize the loop interactions present in the *Fbn2* locus^54,80,81^. We identified nine key loops, with a total of seven anchors comprised of a mix of CTCF binding sites and RNAPII ChIP-seq peaks (Fig. 2b,c, Table S2). RNAPII-anchored loops, as quantified by Absolute Looping Estimator (AbLE) score^58^, were less sensitive to acute degradation of loop extrusion factors than CTCF-anchored loops (which are nearly completely ablated), in accordance with previous work showing that microcompartments are largely robust to depletion of loop extrusion (Fig. 2c, S3b)^54^. These results suggest a role for elements of transcriptional machinery in reshaping the 3D genome that is not influenced by loop extrusion.

**Figure 2:**
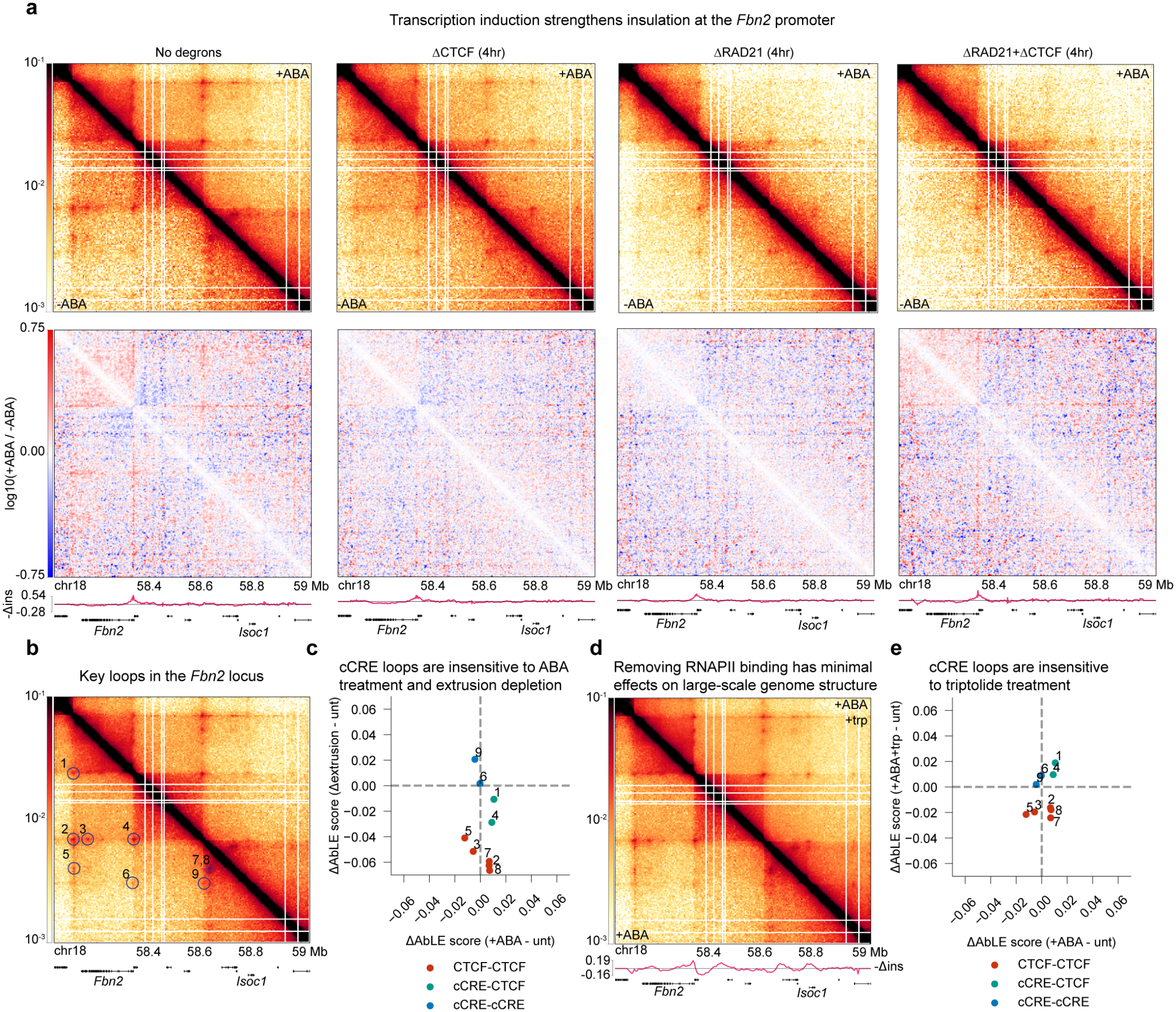
Transcriptional activation creates cohesin-independent insulation across the *Fbn2* promoter. **a**, Contact maps +/-ABA (24h), differential (+ABA/-ABA) contact maps, and differential insulation score tracks (-ABA - +ABA) for P2B2 cells across a range of degron treatments (4h). **b**, Locations of key loops in the *Fbn2* locus indicated on contact map of P2B2 cells +ABA. **c**, Differential AbLE scores (treatment-untreated) for key loops in **b** across transcriptional activation (x-axis) and degradation of RAD21+CTCF (y-axis). **d**, Contact map visualization of RCMC data in P2B2 cells +ABA, +/-triptolide. **e**, As **c**, with ABA+triptolide treatment on the y-axis.

We next aimed to assess whether RNA polymerase II itself plays a role in transcription-mediated 3D genome reorganization as has been suggested^27–30,82^. To remove most RNAPII binding after activating the *Fbn2* promoter, we treated cells with ABA overnight followed by triptolide for 4 hours^54^ (Fig. S3c). Effects of this treatment on contact maps at the *Fbn2* locus compared to ABA alone were minimal, with barrier strength at the *Fbn2* promoter only mildly reduced even after RNAPII depletion (Fig. 2d). RNAPII-anchored loops were also minimally affected by triptolide treatment, consistent with prior work^54,83–85^; however, CTCF-CTCF loops appeared to weaken uniformly upon triptolide treatment (Fig. 2e). Taken in combination with our observation that activating transcription increases promoter insulation even after degradation of loop extrusion factors, these results indicate that direct mechanistic interplay between cohesin and RNAPII (e.g., through blockage of loop extrusion by RNAPII) is not the exclusive mechanism responsible for strengthening insulation at the *Fbn2* promoter. Instead, we propose the following hypotheses, which are not mutually exclusive: that some component or components of transcription machinery recruited by VPR but upstream of RNAPII is responsible for strengthening insulation at the active *Fbn2* promoter, and/or that transcribing RNAPII leaves some mark in the bodies of active genes that contributes to these 3D genome perturbations, potentially by modulating the affinity of the gene body for itself and the surrounding region.

### Transcription increases 3D distances at the *Fbn2* locus and speeds up chromatin dynamics

Cohesin-mediated looping and active transcription are both highly dynamic processes; thus, to more fully characterize their interplay in time as well as space, we turned to super-resolution live-cell imaging^15–17,53^. We tracked the positions of both *Fbn2* TAD anchor CTCF sites and the presence of nascent *Fbn2* MS2-labeled transcripts in P2B2 cells at 30s resolution for up to 3 hours using lattice light-sheet microscopy (Fig. 3a). We extracted inter-TAD-anchor 3D distances and MS2 signal from the 3-color imaging data as previously described^17^. To directly examine the effects of active transcription, we imaged cells both with and without overnight ABA treatment, as well as cells treated with ABA overnight and triptolide for 4 hours. We also imaged cells treated with dTAG-13 or indole-3-acetic acid (IAA) (to degrade RAD21 or CTCF, respectively) for 4 hours to compare changes from transcriptional activation with those from perturbing loop extrusion (Methods). We employed BILD to infer periods of CTCF-CTCF looping^15^ and developed a simple two-state hidden Markov model to segment trajectories into periods of active transcription from MS2 signal (Methods). Examples of the resulting tracks and annotations (from ABA-treated cells) are shown in Figure 3a.

**Figure 3:**
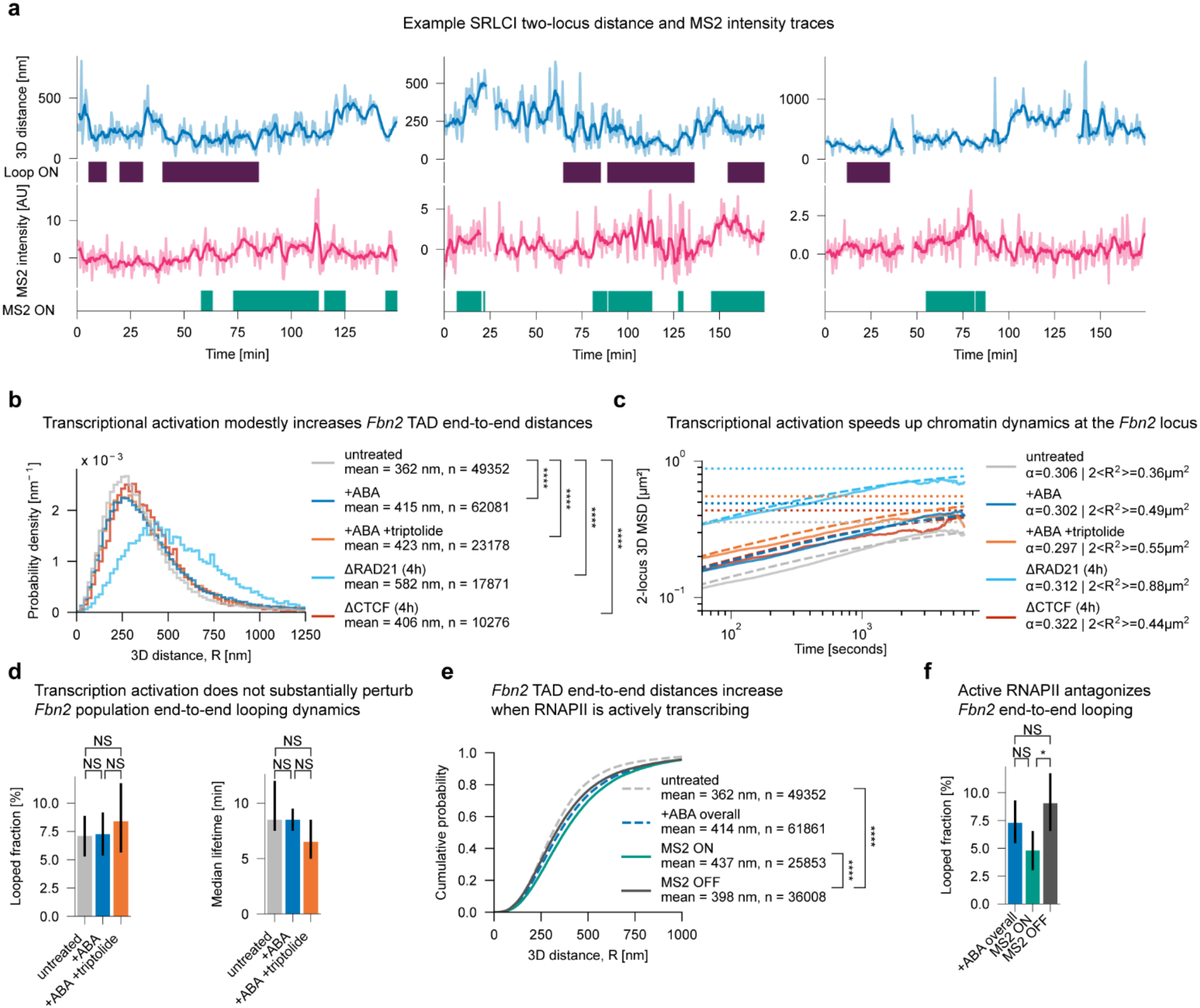
Transcriptional activation speeds up chromatin diffusion, while transcribing RNAPII interferes with cohesin-mediated looping. **a**, Example live-cell 2-point distance and MS2 intensity trajectories, with BILD loop-state annotations and HMM burst annotations. **b**, Probability density distributions of 2-point distances, with mean distances annotated. *P*-values were computed by two-sample K-S test with Bonferroni correction. ****: *P*<0.0001. *P*-values: untreated vs +ABA 2.4e-199, untreated vs +ABA+triptolide 2.7e-112, untreated vs ΔRAD21 <2.2e-308, untreated vs ΔCTCF 4.3e-78. **c**, Localization-error corrected experimental 2-point MSDs (solid lines), analytical MSD fits (dashed lines), and analytical MSD asymptotes (dotted lines). **d**, Mean fully-looped probability and loop lifetime across conditions. Error bars indicate 95% confidence intervals, determined by bootstrapping. *P*-values computed by Welch’s *t*-test with Bonferroni correction. NS: not significant. **e**, Cumulative distribution function of 2-point 3D distances. *P* values computed by 2-sample K-S test with Bonferroni correction. ****: *P*<0.0001. MS2 ON vs MS2 OFF: *P* = 6.91e-107. MS2 OFF vs UNT: *P* = 4.85e-60. **f**, Mean fully-looped probability, stratified by RNAPII state. Error bars represent 95% confidence intervals. *P*-values computed by Welch’s *t*-test with Bonferroni correction. *: *P* < 0.05, NS: not significant. MS2 ON vs MS2 OFF: *P* = 0.027.

We began by examining the distributions of inter-TAD-anchor 3D distances in our live-cell trajectories. We found that cells treated with ABA exhibited a higher mean 3D distance than untreated cells, and that the distribution of distances showed a larger right tail consistent with transcription-induced decompaction^50^ (Fig. 3b). Strikingly, the 3D distance increase from ABA treatment exceeded that of CTCF depletion (Fig. 3b), indicating that transcriptional activation can act as a potent regulator of 3D genome structure on par with CTCF. We also note that 3D distances in untreated cells matched well with C36 cells studied by Gabriele *et al*^15^ (Fig. S4a). We then computed the localization-error-corrected 2-point mean square displacement (MSD)^15,17^; all conditions showed highly subdiffusive dynamics (α≈0.3) (Fig. 3c, Table S3), consistent with recent findings of sub-Rouse dynamics in mESCs^86^. Prior studies have shown contradictory effects of transcription on chromatin dynamics; some report a decrease in chromatin motion upon transcriptional activation that can be reversed by transcription inhibition by drugs such as triptolide^60,87–90^, while another shows faster dynamics upon activation^91^, and yet others report both effects depending on local genomic context^48,92^. However, it has not previously been possible to simultaneously observe chromatin dynamics and inducible nascent transcription at the same locus. Compared to untreated cells, ABA-treated cells displayed faster diffusion, indicated by a higher MSD prefactor and asymptote (Fig. 3c, Table S3). These results are consistent with a model whereby activating transcription leads to decompaction of the locus. All the aforementioned effects were recapitulated in cells treated with ABA followed by triptolide, indicating that the decompaction of the locus can persist after RNAPII leaves the active gene (Fig. 3b,c).

### Active RNAPII is associated with longer end-to-end distances and reduced CTCF-CTCF looping

To investigate possible direct interactions between transcriptional machinery and cohesin, we examined the dynamics of the fully-looped state of the *Fbn2* TAD (i.e., when the CTCF sites are directly connected by one or more cohesin loops), as inferred by BILD^15^. On the level of the entire population, including cells with and without RNAPII currently transcribing, ABA treatment had negligible effect on looping dynamics: both the fraction of fully looped alleles (untreated 7.1%, ABA-treated 7.3%) and the median loop lifetime (untreated 8.5min; ABA-treated 8.5min) were not significantly altered (Fig. 3d), and loops decayed at roughly the same rates (Fig. S4b). However, based on the larger right tail in the +ABA 3D distance distribution, we hypothesized that a sub-population of actively bursting alleles could exhibit more dramatic 3D genome perturbations while the overall population effect remains muted. To test this hypothesis, we computed the distance distributions for active transcription ON (visible MS2 nascent RNA) and active transcription OFF points separately. Actively transcribing alleles exhibited a significant rightward shift in distance distribution, implying that the presence of transcription machinery impacts the 3D genome in addition to the decompaction previously described (Fig. 3e, S4c). We also note that even inactive alleles in the +ABA condition display larger end-to-end distances than untreated cells, indicating that transcription leaves a lasting effect on 3D genome dynamics at the locus (Fig. 3e). Repeating looped fraction estimation on data stratified by MS2 signal revealed that alleles with RNAPII present show significantly reduced looped fraction compared to alleles not actively bursting (RNAPII ON 4.8%; RNAPII OFF 9.1%) (Fig. 3f). These findings provide the first direct live-cell observations consistent with the model that RNAPII may serve as a barrier to cohesin-mediated loop extrusion,^28,29,31,93^ which had previously been primarily inferred from indirect evidence. Overall, we find that nascent transcription results in reduced CTCF-CTCF loop strength, in addition to the more long-lasting, cohesin-independent insulation associated with promoter activation described above.

### Polymer simulations indicate activating transcription induces local decompaction and that polymerase is a barrier to loop extrusion

To understand the mechanistic basis for our observations, we conducted 3D polymer simulations of plausible parameterized mechanistic models of the *Fbn2* region. A high-confidence model would need to reproduce the following observations that we made above: (a) an increase in insulation strength and decrease in 3D contacts across the *Fbn2* promoter, including without loop extrusion factors and/or RNAPII present; (b) relative insensitivity of key CTCF-CTCF loops to transcriptional activation at a whole-population level; (c) an increase in 3D distances between the *Fbn2* TAD anchors; (d) an upward shift of 2-point MSD without altering anomalous exponent; and (e) further increases in 3D distance, and a reduction in looped fraction of the *Fbn2* TAD, when RNAPII is actively transcribing.

We modeled the locus as a series of 2004 coarse-grained monomers representing 500bp of chromatin each^15,28,58,80^. All simulations contained cohesins (modeled as bonds between pairs of monomers that processed bidirectionally along chromatin), CTCF sites that stalled progression of cohesins, and genes from which RNAPII could load and progress along chromatin (Fig. 4a, Methods). To maximize the visibility of moving-barrier effects, genes were constitutively active in these simulations. We tested four models: 1) local decompaction via loss of self-affinity of monomers in the *Fbn2* gene body and nearby region (Fig. 4b); 2) increased self-affinity of monomers in the *Fbn2* gene body and nearby region (Fig. 4c); 3) strengthening of affinity-mediated microcompartmental interactions at RNAPII-bound sites (Fig. 4d); 4) and blockage of cohesin by RNAPII (moving barrier activity) (Fig. 4e, Table S4, Fig. S5-9). We reasoned that local affinity-based decompaction could induce insulation, 3D distance, and MSD increases without major effects on CTCF-CTCF looping (observations a, b, c, d). While we thought increased self-affinity would be less likely to reproduce experimental effects on MSD and 3D distance, we reasoned that it could still generate insulation and 3D contact changes that matched experiment (observations a, b). Since the R1 anchor exhibits a very small amount of RNAPII ChIP-seq signal and the L1 anchor does not, we reasoned that sequestration of the R1 anchor by the *Fbn2* promoter in a microcompartmental-like interaction^54^ could explain the increases in insulation and 3D distance (observations a, c). Finally, we reasoned that RNAPII acting as a moving barrier to loop extrusion would be able to reproduce most experimental 3D genome effects^28^ and would likely also reproduce the increase in 3D distance and decrease in looped fraction during active bursts (observations a, c, d, e). To better understand trends in the simulation results, we simulated a weak, moderate, and strong parameter set for each model.

**Figure 4:**
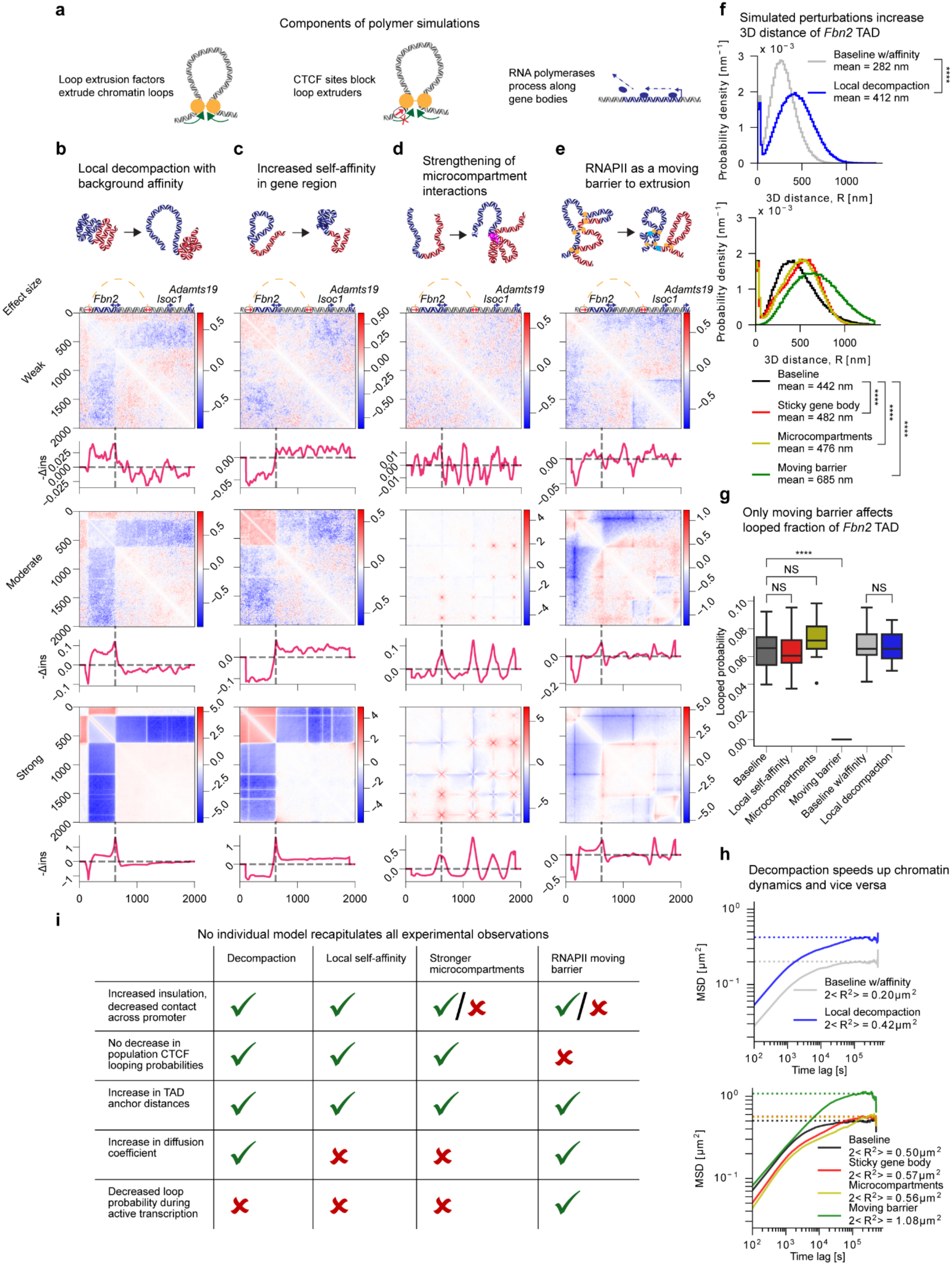
Polymer simulations predict signatures of mechanistic models of transcription-3D genome interaction. **a**, Schematic representation of shared components of all polymer simulations conducted in this work. **b-e**, Schematic representations of simulated perturbations, with differential contact maps (perturbation/baseline) and differential insulation score tracks (baseline - perturbation) for weak, medium, and strong parameters for each model. Blue DNA = gene region; red DNA = rest of locus; magenta DNA = microcompartment anchor. **f**, Probability density distributions of 2-point 3D distances from the strongest simulated parameter sets. *P*-values computed by 2-sample K-S test with Bonferroni correction. ****: *P* < 0.0001. All *P*-values in this panel <2.2e-308. **g**, Fully-looped probabilities for individual simulated repeats of the *Fbn2* TAD. *P*-values computed by Welch’s *t*-test with Bonferroni correction. ****: *P* < 0.0001, NS: not significant. Baseline vs moving barrier *P* = 1.1e-14. **h**, Empirical 2-point MSDs (solid lines) and asymptotes (dotted lines) for simulated perturbations. **i**, Table cataloguing which key experimental observations are satisfied by each simulated model.

Of the models tested, local decompaction (Fig. 4b), increased self-affinity (Fig. 4c), and weak moving-barrier activity (Fig. 4e) produced a strong decrease in interactions between the left and right halves of the *Fbn2* TAD, resembling the effect seen in P2B2 cells after ABA treatment (Fig. 2a). Strengthening microcompartments did not have widespread effects on the locus, while stronger moving barrier models created strong contact depletion across the gene body that was not observed in experimental maps (Fig. 2a, Fig. 4d,e, Fig. S5). With sufficient parameter strength, all models tested reproduced the peak in insulation strength at the *Fbn2* promoter we observed in experimental RCMC (Fig. 2a, Fig. 4b-e). All models tested resulted in a rightward shift of TAD anchor 3D distances compared to baseline (Fig. 4f). However, only the RNAPII moving barrier model could reproduce the experimentally observed reduction in looping probability of the *Fbn2* TAD (Fig 3f), with all other perturbations having no significant effect on looped fraction (Fig. 4g). We note that strong moving barrier parameters also created accumulation of cohesins at the starts and ends of genes (Fig. S9c), which provides a plausible explanation for previous observations of cohesin accumulation at genes that were thought to be due to targeted cohesin loading^37–40,83^. Finally, we examined the dynamic impacts of our models by computing 2-point MSDs between the ends of the *Fbn2* TAD. Local decompaction and moving barrier models both increased MSD compared to baseline, consistent with experimental results; sticky gene and microcompartment models lowered the MSD (Fig. 4h).

Altogether, no single model perfectly explained all experimental observations (Fig. 4i). The local decompaction and moving barrier models came closest, with local decompaction unable to reproduce only the decrease in looped fraction during RNAPII elongation and moving barrier unable to reproduce 3D genomic effects in the absence of loop extrusion factors. Increasing self-affinity could reproduce effects on contact maps, insulation, and 3D distance, but failed to match the increase in diffusion speed and decrease in looped fraction seen in live-cell data. Strengthening microcompartment interactions similarly failed to recapitulate diffusion and looped fraction effects and did not produce contact maps consistent with experimental results. Thus, we next explored a combination of the local decompaction and moving barrier models to see if each component could compensate for the shortcoming of the other.

### Transcription has cohesin-dependent and cohesin-independent effects on the 3D genome

To maximize correspondence with all our experimental observations, we propose a combined mechanistic model wherein active transcriptional machinery interferes with cohesin-mediated loop extrusion while inducing local decompaction at an active locus; when the promoter turns off and transcriptional machinery leaves the locus, the impediment to cohesin activity is removed, but some lasting mark is left that maintains *Fbn2* gene body decompaction. To better match biological reality, we implemented transcriptional bursting at promoters when simulating this model. We used a simplified 1D polymer simulation to optimize values of promoter *k*_*on*_ and RNAPII permeability to cohesin to best match experimental readouts (observed frequency of RNAPII in the *Fbn2* gene body and looped fraction with/without RNAPII, respectively) (Methods, Fig. S10). We selected a value for ε_B_, which determines baseline affinity and thus decompaction strength, to best match the observed mean 3D distance of the *Fbn2* TAD in untreated P2B2 cells (Methods, Fig. S10). 3D polymer simulations of the resulting model faithfully reproduced experimental contact map and insulation patterns (Fig. 2a, Fig. 5a). Simulated contact maps were also robust to depletion of loop extrusion components, in accordance with experimental results (Fig. 2a, Fig. 5b). The joint simulation model reproduced the rightward shift in 3D distance distribution and increase in MSD prefactor seen in live-cell data, with decompaction of the *Fbn2* region as the primary driver of these effects (Fig. 3b,c, Fig. 5c,d). The looped fraction of the *Fbn2* TAD was unaffected on a population level, in line with experimental results (Fig. 3d, Fig. 5e). When RNAPII was present in the simulated gene body, looping probability decreased and 3D distance increased compared to when RNAPII was not actively transcribing, also in line with experimental observations (Fig. 3f, Fig. 5f,g). This joint 3D simulation thus points to a combined model in which active transcription reshapes the 3D genome in both a cohesin-dependent manner by impeding loop extrusion and a cohesin-independent manner by decompacting the active locus.

**Figure 5:**
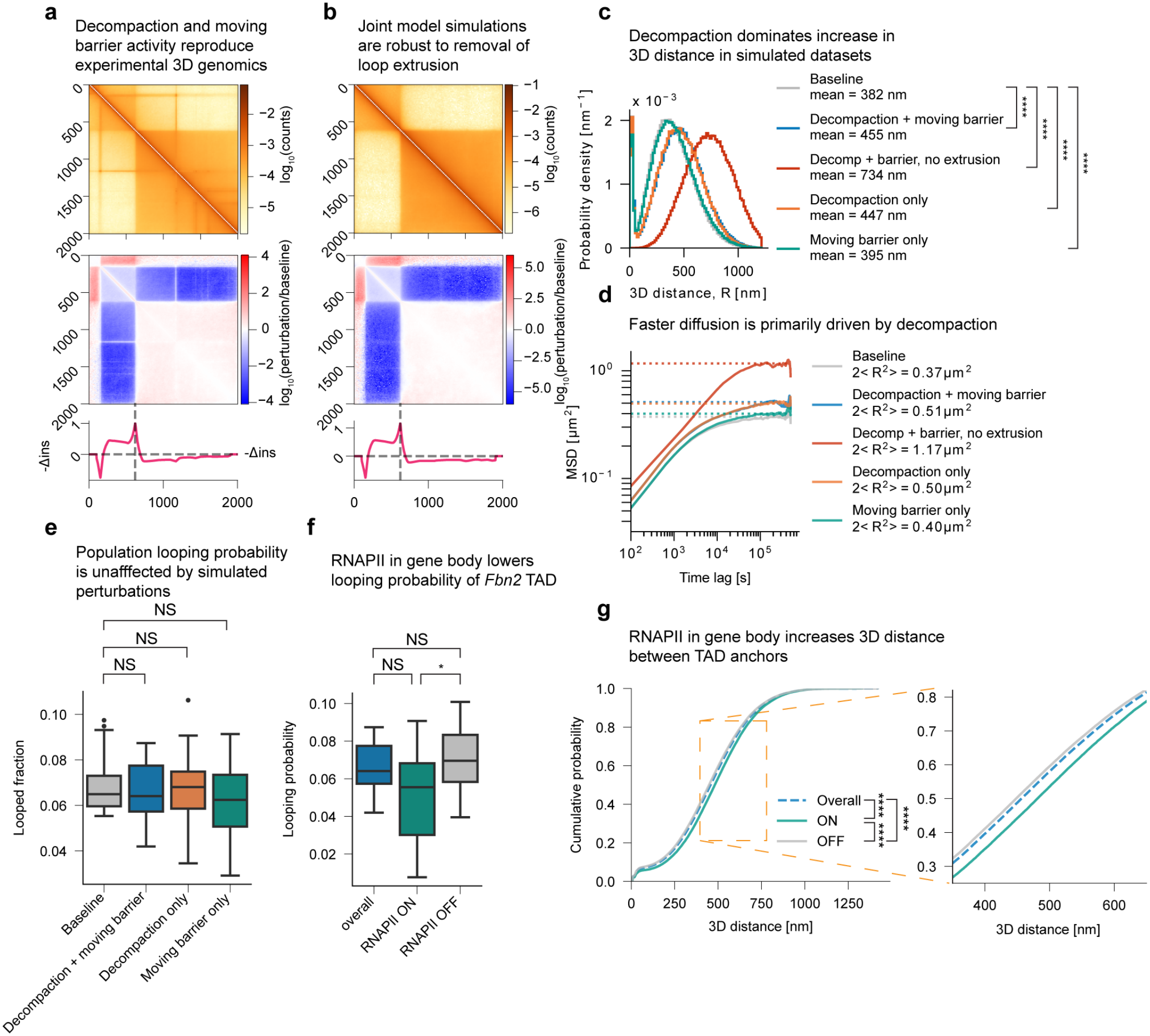
Local decompaction and RNAPII extrusion barrier activity combined can recapitulate all experimental observations. **a**, Raw and differential (perturbation/baseline) contact maps and differential insulation score track (baseline - perturbation) for a combined simulation incorporating both local decompaction and RNAPII barrier activity. **b**, As **a**, for simulations with LEFs removed. **c**, Probability density distributions of 2-point 3D distances for combined simulation model and perturbations removing individual components. *P*-values computed by 2-sample K-S test with Bonferroni correction. ****: *P* < 0.0001. *P*(baseline vs moving barrier only) = 2.3e-63; all other *P*-values in this panel <2.2e-308. **d**, Empirical 2-point MSDs (solid lines) and asymptotes (dashed lines) for combined simulation model and perturbations removing individual components. **e**, Fully-looped probability for individual simulated repeats of the *Fbn2* TAD across different simulation conditions. *P*-values computed by Welch’s *t*-test with Bonferroni correction. NS: not significant. **f**, As **e**, for combined simulation model only, stratified by RNAPII activity in *Fbn2* gene body. *P*-values computed by Welch’s *t*-test with Bonferroni correction. *: P < 0.05, NS: not significant. RNAPII ON vs OFF: *P* = 0.024. **g**, Cumulative distribution function of 2-point 3D distances for combined simulation model, stratified by RNAPII presence. *P*-values computed by 2-sample K-S test with Bonferroni correction. ****: *P* < 0.0001. *P*-values: overall vs ON 3.0e-77, overall vs OFF 1.2e-18, ON vs OFF 1.8e-129.

## DISCUSSION

In this work, we comprehensively characterize the spatial and temporal interaction between cohesin-mediated loop extrusion and active transcription by integrating 3D genomics, super-resolution live-cell imaging, and polymer simulations. Our findings reconcile previous studies that suggest that transcription acts on the 3D genome either via direct interference with loop extrusion^28–35^ or by cohesin-independent modulation of contacts in and around active genes^31,49,50,94^ and support a model whereby transcription acts both in cohesin-dependent and cohesin-independent manners simultaneously (Fig. 5, 6).

**Figure 6:**
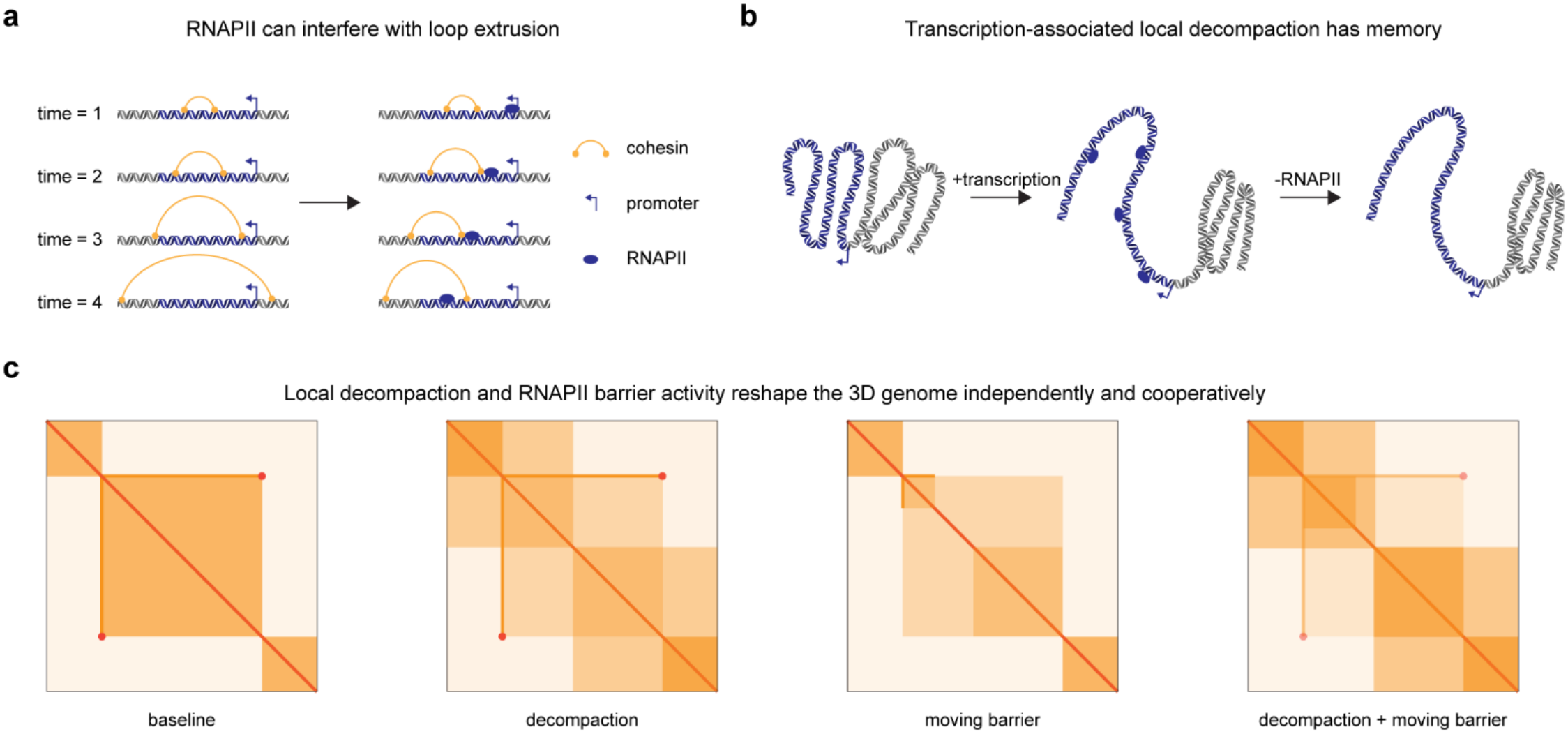
Key takeaways from the combined mechanistic model. **a**, Schematic representation of RNAPII interfering with loop extrusion by blocking cohesins. **b**, Schematic representation of local gene decompaction upon transcriptional activation being retained after transcribing RNAPII is removed. **c**. appearance in 3D maps.

First, our studies support a cohesin-dependent effect of transcription. Specifically, we provide evidence in support of a moving-extrusion-barrier function of RNAPII, which had previously been inferred primarily from simulation data^28,29,31,93^, for the first time via live-cell imaging (Fig. 6a). Our observations of decreased cohesin loop probability and increased 3D distance during active RNAPII elongation (Fig. 3) support that RNAPII can block and reposition cohesins^28,29^ (Fig. 4).

Second, our studies support a cohesin-independent effect of transcription. We find that transcriptional activation strengthens an insulation boundary at the Fbn2 promoter, which splits the wild-type 505-kb TAD into two smaller “TAD-like” domains, and that this insulation boundary persists even without loop extrusion factors present (Fig. 2). This necessitates a cohesin-independent function. The increased diffusion coefficient of the *Fbn2* locus in our live-cell imaging (Fig. 3c), in conjunction with simulation data (Fig. 4h), suggests that this effect is gene-body decompaction^31,49,50^. Integrating these cohesin-dependent and -independent mechanisms, our polymer simulations suggest that at the *Fbn2* locus, cohesin-independent gene body decompaction is the dominant driver of promoter-associated insulation upon transcriptional activation, with the barrier activity of RNAPII playing a lesser role at the levels of transcription observed in our model system (Fig. 5). We also show that gene body decompaction persists even after RNAPII leaves the active locus (Fig. 6b).

Our observation of strong cohesin-independent insulation boundary formation may explain several puzzling observations in the field. First, a major conundrum in the field has been the observation that the expression of some genes is strongly reliant on cohesin-mediated loop extrusion^17,25,95–97^, whereas the global transcriptional effects of cohesin depletion are more modest^12,98,99^. In general, gene-rich genomic regions tend to be more robust to cohesin-depletion. Insulation boundary formation at active promoters^6,11^ through transcription-associated gene-body decompaction may help explain why gene expression is relatively robust to cohesin depletion. By preserving promoter insulation independently of loop extrusion, this mechanism could limit inappropriate enhancer–promoter interactions and reduce transcriptional dysregulation following cohesin loss.

Second, our findings may provide a mechanistic explanation for promoter competition^100–103^. Promoter competition is supported by strong genetic evidence, but the mechanistic basis has remained mysterious. A recent study by Koska *et al*. systematically studied promoter competition by inserting reporter genes between the endogenous *Sox2* gene and enhancers. Koska *et al*. demonstrated that promoter competition depended on transcription and scaled with insulation and transcript length, but surprisingly was cohesin-independent^103^. Our results here suggest that transcription-induced gene body decompaction leading to cohesin-independent insulation boundary formation provides a plausible mechanistic explanation for the observations in Koska *et al*.^103^ and perhaps promoter competition more broadly^100–103^.

A limitation of our work is the focus on a single engineered locus. Future studies will be required to test the generality of our findings at other loci and in other cell types. Nevertheless, our study emphasizes the power of integrating live-cell imaging and 3D genomics with 3D polymer modeling, which is only possible at single loci. Neither live-cell imaging nor 3D genomics alone would have been sufficient to distinguish all mechanistic models above (Fig. 4-5). In summary, the principles of transcription-mediated decompaction and RNAPII extrusion barrier activity we outline here provide a more complete view of the structural impact of active genes than previously available and suggest a more complex – and less cohesin-centric – view of 3D genome regulation.

## Acknowledgements

The authors thank Ed Banigan for insightful discussions and for sharing code. We thank Luke Lavis for providing Janelia Fluor dyes. We thank Stanley Qi for providing a PYL1/ABI plasmid^104^ and George Church for providing a VPR plasmid^64^ used to create genome editing constructs used in this work. We thank Miles Huseyin, Clarice Hong, Sarah Nemsick, Ed Banigan, Sim Sakong, Sumin Kim, and Varshini Ramanathan for providing critical feedback on the manuscript. We further thank the members of the Hansen lab for constructive discussions and support during the project. We thank the Flow Cytometry Core and MIT BioMicro Center of the MIT Koch Institute’s Robert A. Swanson (1969) Biotechnology Center for technical support. We also thank the Walk-up Sequencing services of the Broad Institute of MIT and Harvard for Illumina sequencing. We thank the High-Content Imaging Facility team at the Center for Development of Therapeutics, Broad Institute, for use and maintenance of their Opera Phenix High-Content Screening System. We thank Novogene USA for RNA-seq services.

## Author Contributions

A.S.H. and J.A.D. designed the project. J.A.D., A.J., and A.S.H. generated and validated the P2B2 cell line. J.A.D. and S.M. conducted and analyzed Micro-C and Region-Capture Micro-C. J.A.D. performed and analyzed all other genomics experiments. J.A.D. performed live-cell imaging. J.T. optimized the Fyrtarn pipeline for the data in this study. J.A.D. and J.T. performed live-cell movie processing, and J.A.D. analyzed the resulting outputs. J.A.D. performed and analyzed all other experiments. J.A.D. performed and analyzed 3D polymer simulations. J.A.D. drafted the manuscript and generated the figures, with input from all authors. A.S.H. supervised all research. J.A.D. and A.S.H. edited the manuscript and figures, with input from all authors.

## Data Availability

All genomics data generated in this study are publicly available on the NCBI Gene Expression Omnibus (GEO) in SuperSeries GSE355306. Previously published Micro-C and RCMC data are available on GEO as follows: Nora *et al*. GSE98671, Bonev *et al*. GSE96107, Hsieh *et al*. GSE130275, Gabriele *et al*. GSE187487, Goel *et al*. GSE207225, Narducci and Hansen GSE256246, Jusuf *et al*. GSE286495. Previously published ChIP-seq data are available on GEO as follows: CTCF and RAD21 GSE90994, RNAPII GSE58019. 3D distance and MS2 intensity trajectories extracted from live-cell movies are available at https://doi.org/10.5281/zenodo.24208285.

## Code Availability

All new code and scripts described in this study are available at https://github.com/ahansenlab/transcription_3D_genome_code/.

## AI Disclosure

No LLMs were used in the production of this manuscript.

