## Supplementary material for "Transcriptional activation forms insulation boundaries through cohesin-independent decompaction and cohesin-dependent barrier formation": Methods and Supplementary Figures and Tables

[1] Department of Biological Engineering, Massachusetts Institute of Technology; Cambridge, MA 02139, USA, [2] Gene Regulation Observatory, Broad Institute of MIT and Harvard; Cambridge, MA 02142, USA, [3] The Novo Nordisk Foundation Center for Genomic Mechanisms of Disease, Broad Institute of MIT and Harvard; Cambridge, MA 02142, USA, [4] Koch Institute for Integrative Cancer Research; Cambridge, MA 02139, USA, [5] Department of Cell Biology, Yale School of Medicine; New Haven, CT 06520, USA, [6] Program in Computational and Systems Biology, Massachusetts Institute of Technology; Cambridge, MA 02139, USA, [7] Department of Mathematics, Howard University; Washington, DC 20059, USA

#### Contents

- Methods
- Methods References
- Supplementary Figures and Figure Legends
- Supplementary Tables

#### METHODS

##### Cell culture

JM8.N4 mouse embryonic stem cells[1] were cultured in feeder-free conditions at 37°C and 5.5% $CO_2$, in KnockOut DMEM (ThermoFisher 10829-018) supplemented with 15% fetal bovine serum (FBS, Avantor 89510-186, lot 266M25), 1000U/mL leukemia inhibitory factor (LIF, Cell Guidance Systems GFM200), 1X GlutaMAX supplement (ThermoFisher 35050-061), 1X MEM Non-Essential Amino Acid Solution (ThermoFisher 11140-050), 0.1mM β-mercaptoethanol (Sigma M-3148), 100µg/mL penicillin-streptomycin (ThermoFisher 15140-122), and 2i: 1µM MEK inhibitor (Tocris PD0325901) and 3µM GSK inhibitor (Sigma SML1046-25MG). Plates and dishes were pre-coated with 0.1% gelatin solution before plating cells. Cells were fed by replacing 50% medium every day and passaged every 2-3 days by incubating with TrypLE Express Enzyme (ThermoFisher Scientific 25300062) for 3-5 minutes at 37°C to detach before resuspending in growth medium, centrifuging at 300*g* for 3 minutes, and re-plating or harvesting.

HEK293T cells (ATCC® CRL-3216™) for spike-in controls were cultured at 37 °C with 5.5% $CO_2$ in DMEM +phenol red (ThermoFisher 11965-092) supplemented with 10% FBS (Avantor 89510-186, lot 266M25), 1x MEM non-essential amino acids, 1x penicillin/streptomycin, 1X GlutaMAX, and 50 µM β-mercaptoethanol. For passaging, cells were incubated with TrypLE for 3 minutes at 37 °C, then resuspended in growth medium before passaging to a new plate. Cells were harvested by centrifugation at 300*g* for 3 minutes.

*Drosophila melanogaster* SG4 cells (RRID:CVCL_Z707; https://dgrc.bio.indiana.edu//stock/RRID:CVCL_Z707) for spike-in were cultured at 25°C in Schneider's *Drosophila* medium (ThermoFisher 21720024) supplemented with 10% FBS (Avantor 89510-186, lot 266M25) and 100µg/mL penicillin/streptomycin. Prior to use, FBS was heat inactivated at 56°C for 30 minutes, then frozen at -20°C and thawed before use. SG4 cells were detached by pipetting for passaging and harvesting. For passaging, cells were plated in a 1:5 ratio of cultured to fresh medium. For harvesting, cells were centrifuged at 500*g* for 5 minutes.

##### Genome editing

Cas9-mediated genome editing was performed in JM8.N4 mESCs according to published procedures[2] with modifications. Cells were grown as previously described to approximately 80% confluency on a 6-well plate or 10cm dish prior to transfection. For each edit, a Cas9 plasmid (encoding Cas9, mVenus, and one or two sgRNAs chosen for each edit and inserted by restriction ligation) and an HDR plasmid (containing the DNA sequence to be inserted and 500-1000bp of homology on the left and right sides) were co-transfected at a 1:4 ratio (500ng Cas9 vector to 2µg HDR vector per well of a 6-well plate, or 3µg Cas9 vector to 12µg HDR vector per 10cm plate) using Lipofectamine 3000 (ThermoFisher L3000015) following manufacturer's protocol. sgRNAs were designed using the CRISPOR tool[3] and cloned into a Cas9 delivery plasmid via restriction ligation. Cas9+sgRNA plasmids were transfected alongside the HDR plasmid. The day after transfection (16-28 hours later), cells were collected via TrypLE and sorted for YFP fluorescence against an untransfected control by FACS. Fluorescent (successfully transfected) cells were plated at a density of 50,000 per 15cm dish. Cells were allowed to grow into colonies for 7 days, at which time single colonies were picked under a Lynx EVO stereomicroscope (Vision Engineering) using a 20µL pipette tip and passaged into 96-well plates. After 3-4 days of expansion and splitting into two 96-well plates, cells from one plate were collected, resuspended in squishing buffer (10mM Tris-HCl pH 8 (Thermo Fisher 15568025), 1mM EDTA (Invitrogen 15575020), 25mM NaCl (Millipore Sigma S6546), 200µg/mL Proteinase K (Viagen 501-PK)), and lysed at 65°C for 1 hour and 95°C for 10 minutes to collect crude genomic DNA. Clones were screened for successful edits by PCR. Clones that passed initial screening were expanded further; clean gDNA was extracted with a Zymo Quick-DNA kit (Zymo Research D3024) and edits were validated by PCR screening and Sanger sequencing (Azenta). See Table S5 for plasmids and sgRNA sequences and Table S6 for primer sequences.

Our previously published cell lines[4] for tracking the CTCF loop anchors of the *Fbn2* TAD were derived from mESC clone C59 where both CTCF and cohesin are tagged (FLAG-Halo-CTCF and RAD21-SNAP$_f$-V5)[5]. Here, we needed to use the Halo channel for RNA visualization. We therefore started from wild-type JM8.N4 mouse embryonic stem cells[1] rather than working from the previously published cell lines in Gabriele *et al.*[4]. In wild-type mESCs, we began by tagging the CTCF loop anchors using the same genome-editing constructs reported in Gabriele *et al.*[4]. We used Cas9-mediated genome editing as described above to insert a 224xTetO array ~5kb upstream of the L1 CTCF site and an ANCH3 (NeoVirTech) array ~5kb downstream of the R1 site, following the targeting strategy of Gabriele *et al.*[4], to produce clone C69. To improve localization precision in live-cell imaging, we next aimed to shorten the tether lengths between the arrays and the CTCF sites by deleting ~4kb of the intervening sequence. Two serial genome edits were performed as described above to delete the L1 tether region and R1 tether region; no repair plasmid was used for these deletions. After PCR screening and Sanger sequencing validation, three clones (P4A11, P4E6, and P4E9) were validated to have a deletion corresponding to chr18:58,104,237-58,109,227 of the mm39 wild-type genome build, resulting in a tether length of 946bp on the L1 side. The R1 deletion was performed on all three clones; after screening, one clone (P2D9) was validated to have a deletion corresponding to chr18:58,615,476-58,618,792 (mm39) resulting in a tether length of 897bp on the R1 side.

Next, we aimed to tag *Rad21* with a degron tag to allow acute degradation. We used Cas9-mediated genome editing as described above to insert FKBP12$^{F36V}$ at the C-terminus of *Rad21* (mm39 chr15:51,827,732) and produce clone P1G9. We then sought to insert an inducible activation system to allow control of transcriptional activation of *Fbn2*. To achieve this, we used Cas9-mediated editing as previously described to insert PYL1-VPR[6,7] and ABI-LacI[6] constructs at the *Hipp11* safe-harbor locus, following the targeting strategy employed by Browning *et al*[8]. Two clones, P2F12 and P4C3, were validated to contain both constructs at chr11:3,195,344 by PCR and Sanger sequencing. After this, we aimed to tag *Ctcf* with an orthogonal degron tag to allow acute degradation of *Ctcf* independently of *Rad21*. We inserted mAID at the N-terminus of *Ctcf* (mm39 chr8:106,390,401) to produce clone P5G7. Next, to allow transcriptional activation of *Fbn2* and visualization of nascent transcripts, we simultaneously inserted a 6xLacO array into the *Fbn2* promoter (mm39 chr18:58,343,305) and a 126xMS2 array into the first exon of *Fbn2* (mm39 chr18:58,342,737). One clone, P1C3, was validated to contain both arrays.

The AID system requires cells to express *Oryza sativa* TIR1 (OsTIR1) to induce degradation of the target protein[9]; we introduced this protein via PiggyBac transposase-mediated random integration. Cells were grown to ~80% confluency in one well of a 6-well dish, then transfected with 2.5μg of a plasmid encoding OsTIR1 and a blasticidin resistance marker (pASH133) using Lipofectamine 3000 following manufacturer's protocol. After transfection, cells were grown in medium containing 10μg/mL blasticidin (Sigma-Aldrich 15205) for five days (until all cells in an untransfected control dish had died) to select for successfully transfected cells.

Finally, to enable tracking of the positions of *Fbn2* TAD anchors and transcription of *Fbn2*, we randomly integrated tdMCP-2x-HaloTag[10–13], TetR-2x-mScarlet3[14] (pASH424), and OR3-2x-m(n1)oxStayGold[13,15,16] using PiggyBac transposase as described above. One day after transfection, cells were stained with Halo-JFX650 dye (Promega HT1070) and sorted for GFP, RFP, and JFX650 fluorescence by FACS against an untransfected control. To select for optimal expression levels of the fluorophores, fluorescent cells were subcloned onto optically-clear 96-well plates (Revvity 6055302) and imaged on an Opera Phenix microscope (Revvity). Clone P2B2 was selected for optimal signal/noise ratio of the dots corresponding to DNA loci and nascent transcription bursts.

##### Activation of transcription at *Fbn2* locus

To activate transcription of *Fbn2*, P2B2 cells were cultured in medium containing 100μM abscisic acid (ABA) overnight (16-24 hours). Activation of *Fbn2* was confirmed by RT-qPCR and RNA-seq. For RT-qPCR, cells were treated with ABA overnight, then collected via TrypLE. RNA was extracted using a Zymo Quick-RNA kit (Zymo Research R1057) following manufacturer's instructions. RT-qPCR was conducted using a LightCycler 96 (Roche) and a Luna RT-qPCR kit (NEB E3005) following manufacturer's instructions (see Table S6 for qPCR primer sequences). RNA-seq was performed by Novogene.

##### Acute degradation of RAD21 and CTCF

RAD21-FKBP12$^{F36V}$ was acutely degraded by adding dTag-13 (Fisher Scientific 66-055) to the medium at a final concentration of 500nM for 4 hours. mAID-CTCF was acutely degraded by adding indole-3-acetic acid (IAA, Sigma-Aldrich I5148) at a final concentration of 500μM to the medium for 4 hours. Degradation was validated by Western blot, following the protocol of Cattoglio *et al.*[17] with modification. Cells were grown in 10cm plates, harvested with TrypLE, washed once with PBS (Thermo Scientific 10010049), and lysed with high-salt lysis buffer (500mM NaCl, 25mM HEPES (Teknova H1035), 1mM $MgCl_2$ (Millipore Sigma M9272), 0.2mM EDTA, 0.5% IGEPAL CA-630 (Sigma-Aldrich I8896), 1X PMSF (ApexBio A2587)). 4X Laemmli Sample Buffer (Bio-Rad 1610747) was added to a final concentration of 1X and samples were boiled at 95°C for 15 minutes.

Western blotting was performed using Novex™ 4-12% Tris-Glycine Mini Protein Gels (ThermoFisher XP04122) in premade Tris/glycine/SDS running buffer (Fisher Scientific LC26754). Gels were loaded with samples and Bio-Rad Precision Plus Protein™ Dual Color Standards (Bio-Rad 1610394) and run at 200V for 45 minutes. Proteins were transferred to pre-equilibrated nitrocellulose membranes (Bio-Rad 1620112) using a GenScript eBlot™ L1 wet transfer machine (GenScript L00686). Membranes were blocked for 30 minutes with 5% nonfat milk (Genesee Scientific 20-241) in TBS with 0.1% Tween-20 (VWR M147) (TBS-T). Primary immunostaining was conducted overnight at 4°C in 5% nonfat milk in TBS-T with the following antibodies at 1:1000 dilution: Abcam ab992 (RAD21), Millipore Sigma 07-729 (CTCF), Abcam ab818 (TBP). The next day, blots were washed once with TBS-T and three times with TBS, then secondary immunostaining was conducted for 1 hour at room temperature in 5% nonfat milk in TBS-T with the following antibodies at 1:5000 dilution: Rockland 18-8816-31 (anti-rabbit IgG) and Rockland 18-8817-31 (anti-mouse IgG). Blots were washed once with TBS-T and three times with TBS, then treated with Pierce™ ECL Western Blotting Substrate (ThermoFisher 32106) according to manufacturer's instructions and imaged on a ChemiDoc™ imaging system (Bio-Rad 12003153). Protein quantification was performed using Fiji[18].

##### Micro-C and RCMC

For RCMC[19], cells were grown in 15cm dishes, harvested with TrypLE, washed in PBS with 1% bovine serum albumin (BSA, Gibco 15260037), and resuspended in PBS. For each sample, 30M resuspended cells were crosslinked in 30mL PBS with 3mM disuccinimidyl glutarate (DSG, ProteoChem c1104) for 35 minutes, then 16% formaldehyde (Thermo Scientific 28908) was added to a final concentration of 1% v/v and crosslinking continued for 10 minutes. Crosslinking was quenched with Tris pH 7.5 (Fisher Scientific 50-103-1366) at a final concentration of 0.375M for 5 minutes at room temperature and 5 minutes on ice. Crosslinked cells were split into 5M cell aliquots, pelleted in a benchtop microcentrifuge, snap-frozen in liquid nitrogen, and stored at -80°C.

Micro-C and RCMC were performed following the protocol of Huseyin *et al.*[19] with minor modifications. Briefly, the protocol consists of micrococcal nuclease (MNase) titration, chromatin digestion and labeling, proximity ligation, reverse crosslinking, size selection and purification, library preparation, and (for RCMC) target enrichment.

For MNase titration, 5M crosslinked cells per titration were thawed on ice and incubated for 20 minutes in 1mL of buffer MB1 (50mM NaCl, 10mM Tris-HCl pH 7.5 (Fisher Scientific 15-567-027), 5mM $MgCl_2$, 1mM $CaCl_2$ (Millipore Sigma C4901), 0.2% IGEPAL CA-630, 1x cOmplete™ Protease Inhibitor Cocktail (Roche 04693159001)). Cells were centrifuged at 1750*g* for 5 minutes and supernatant removed, washed once with MB1 and centrifuged, then resuspended in 500μL MB1 and split into five 1M cell aliquots. MNase (Worthington Biochemical LS004798) was added to each aliquot at a range of final concentrations (typically 4U-14U per million cells), and the mixtures incubated at 37°C for 20 minutes with shaking at 1000rpm. Digested chromatin was centrifuged at 300*g* and 4°C for 5 minutes and the supernatant removed. Chromatin pellets were resuspended in 150μL of Tris-EDTA buffer (TE, Sigma-Aldrich 93283) with 1mg/mL Proteinase K, 1% sodium dodecyl sulfate (SDS, Sigma-Aldrich L3371), 0.1mg/mL RNase A (Thermo Scientific EN0531), and 200mM NaCl and incubated at 65°C with shaking at 1000rpm overnight to reverse crosslinks. The next day, DNA was purified with a Zymo DNA Clean and Concentrator kit (Zymo D4034) following manufacturer's instructions. Purified DNA was run on a 1% agarose gel at 120V for 1 hour. The MNase concentration with a mononucleosome:dinucleosome ratio closest to 85:15 was selected as optimal.

For RCMC, 25M crosslinked cells (5 tubes of 5M cells each) per replicate were thawed, incubated in MB1, and washed and resuspended as for MNase titration. MNase was added in the optimal concentration as determined by titration and cells incubated at 37°C for 20 minutes with shaking at 1000rpm. Digestion was quenched by adding EGTA (Bioworld 40520008-2) to a final concentration of 4mM, after which nuclei were centrifuged at 1750*g* and 4°C for 5 minutes and supernatant removed. Nuclei were washed twice with 250μL of buffer MB2 (50mM NaCl, 10mM Tris-HCl pH 7.5, 10mM $MgCl_2$, 100μg/mL BSA (Sigma-Aldrich B8667)). Pelleted nuclei were resuspended in 100μL of a solution of 50mM NaCl, 10mM Tris-HCl pH 7.5, 10mM $MgCl_2$, 100μg/mL BSA, 2mM ATP (ThermoFisher R1441), 5mM dithiothreitol (DTT, Sigma-Aldrich 10197777001), and 0.5U/μL T4 PNK (NEB M0201) and incubated at 37°C with shaking at 1000rpm for 15 minutes, followed by the addition of DNA Polymerase I Klenow fragment (NEB M0210) to final concentration of 0.5U/μL and incubation for a further 15 minutes to chew back DNA ends for subsequent biotin labeling. To label chewed ends with biotin, 50μL of a solution containing 200μM each of biotin-14-dATP (Jena Bioscience NU-835-BIO14), biotin-11-dCTP (Jena Bioscience NU-809-BIOX), dGTP (Jena Bioscience NU-1003), and dTTP (Jena Bioscience NU-1004), 100μg/mL BSA, and 1X T4 DNA Ligase Buffer (NEB B0202) was added to each tube of nuclei and the mixtures incubated for 45 minutes at 25°C with interval shaking (1min 1000rpm, 3min off). After biotin labeling, nuclei were centrifuged at 1750*g* and 4°C for 5 minutes and supernatant removed. Pellets were washed once with 250μL of buffer MB3 (50mM NaCl, 10mM $MgCl_2$, 100μg/mL BSA). For proximity ligation, nuclei pellets were resuspended in 500μL of a solution of 1x T4 DNA ligase buffer, 100μg/mL BSA, and 20U/μL T4 DNA ligase (NEB M0202) and incubated with rotation at room temperature overnight.

The following day, ligated mixtures were centrifuged at 3000*g* and 4°C for 5 minutes and supernatant removed. Ligated DNA was resuspended in 200µL of 1X NEBuffer 1 (NEB B7001) with 5U/µL Exonuclease III (NEB M0206) and incubated at 37°C with interval shaking (1min 1000rpm, 3min off) for 15 minutes to remove biotinylated nucleotides from unligated ends. Proteinase K to 1mg/mL, SDS to 1%, RNase A to 0.1mg/mL, and NaCl to 200mM were added to the solution, and the mixture incubated at 65°C overnight with shaking at 1000rpm to reverse crosslinks. The next day, DNA was purified from the reverse crosslinking mixture with a Zymo DNA Clean and Concentrator kit according to manufacturer's instructions, with two elutions of 25µL each instead of one. Purified DNA was run on a 1% agarose gel at 120V for 1 hour, after which the 250-400bp band was cut out with a scalpel blade. Dinucleosome-sized DNA was extracted from the agarose gel with a Zymo Gel DNA Recovery kit (Zymo D4008) following manufacturer's protocol, with two elutions of 15µL each instead of one.

For each 5M cells of input, 50µL Dynabeads MyOne T1 streptavidin beads (Invitrogen 65601) were washed twice in TBW (5mM Tris-HCl pH 7.5, 1M NaCl, 0.5mM EDTA, 0.1% Tween-20) and pelleted on a magnetic rack. After removal of supernatant, beads were resuspended in 150µL 2X BW (10mM Tris-HCl pH 7.5, 2M NaCl, 1mM EDTA), then labeled and size-selected DNA was added and the total volume brought to 300µL per 5M cells of input with nuclease-free water (Fisher Scientific AM9937). The mixture was incubated at room temperature overnight with rotation to allow biotinylated DNA to bind to beads. The next day, beads were collected on a magnetic rack and washed twice with TBW and once with 10mM Tris-HCl pH 7.5, then resuspended in 50µL Zymo elution buffer (Zymo D3004).

Library preparation was conducted using reagents from the NEBNext™ Ultra II DNA Library Prep kit (NEB E7645), following a modified version of the manufacturer's protocol. For each 50µL of beads, 7µL of End Prep Reaction Buffer and 3µL of End Prep Enzyme Mix were added and the mixture incubated at 20°C with interval mixing (1min 1000rpm, 3min off) for 30 minutes, followed by 30 minutes at 65°C with interval mixing as above. To this mixture were added 2.5µL NEBNext™ Unique Dual Index UMI Adaptors (NEB E7395), 30µL Ligation Master Mix, and 1µL Ligation Enhancer. The mixture was incubated at 20°C with interval mixing as above for 15 minutes. Beads were then washed twice on a magnetic rack with TBW and once with 10mM Tris-HCl pH7.5 and resuspended in Zymo elution buffer. As each replicate contained multiple tubes, those tubes were pooled at this step. Libraries were PCR amplified using NEBNext™ Ultra II Q5 Master Mix and Universal Primers following manufacturer's instructions, with target yield of 500ng (typically 6-7 PCR cycles). To purify DNA following PCR, libraries were mixed with AMPure XP beads (Beckman Coulter A63880) to a final concentration of 0.9X and incubated at room temperature for 5 minutes. Beads were pelleted on a magnetic rack and washed twice with freshly-prepared 80% ethanol, then air-dried for 5 minutes on the rack. DNA was eluted from beads by adding Zymo elution buffer and incubating at room temperature for 2 minutes. Beads were pelleted on the magnetic rack and the supernatant containing the finished libraries transferred to a new tube. Target enrichment was performed on purified libraries using a Twist Bioscience Standard Hybridization Reagent Kit (Twist Bioscience 104447) following manufacturer's protocol. A custom probe panel for target enrichment was designed in collaboration with Twist. After target enrichment, libraries were sequenced on an Illumina NovaSeq X.

**RCMC data analysis**

Paired-end reads in .fastq format from sequencing Micro-C/RCMC libraries were aligned to the WT mm39 genome or an edited genome build for clone P2B2, as appropriate, using bwa-mem2[20]. Aligned reads were paired using pairtools[21] parse with --walks-policy set to "expand", and duplicate reads (due to PCR duplication) were removed with pairtools dedup with --max-mismatch set to 1. The resulting .pairs files were indexed with pairix[22]. Pairs files were first filtered for pairs with both ends lying in the captured region using pairtools[21] select, after which the filtered .pairs files were indexed and parsed into contact maps at high resolution (small bin size) in .cool format using cooler[23] cload pairs, and then into multi-resolution .mcool files using cooler zoomify. Standard ICE balancing was performed on .mcool files using the --balance flag of cooler zoomify. Neighbor balancing was performed on .mcool files using the genome-wide protocol of the neighbor-balance package[24].

Raw and differential contact maps were plotted using CoolBox[25]. Stratum-adjusted correlation coefficients between .mcool files were computed using a Python implementation of the hicrep algorithm[26]. Insulation scores were computed as the mean value of a sliding square contact map window with one corner on the diagonal, normalized to the mean of all insulation scores on each map. Loops were annotated by hand on ICE-balanced P2B2 +ABA RCMC data at 5kb resolution. Loop anchors were classified by overlap with CTCF (GSE90994) and RNAPII (GSE58019) ChIP-seq tracks.

**Acute depletion of RNAPII binding and calibrated RNAPII ChIP-seq**

To acutely inhibit RNAPII binding to chromatin, cells were treated with 1µM triptolide for 4 hours before harvesting. Spike-in calibrated RNAPII ChIP-seq[27–29] was performed as previously described[30]. Three replicates were performed for each condition. For each sample, 50M mESCs were grown as previously described on 15cm dishes, harvested with TrypLE, washed with PBS, and resuspended in 10mL PBS. 16% formaldehyde was added to a final concentration of 1% and cells incubated for 10 minutes to crosslink them. Crosslinking was quenched with 1.5mL of 1M glycine for 10 minutes, after which cells were pelleted at 1000*g* and 4°C for 5 minutes, supernatant removed, and cells snap-frozen in liquid nitrogen and stored at -80°C until ready to proceed with the protocol. HEK293T cells for spike-in were grown, crosslinked, and frozen separately as above; 2M HEK293T cells were collected for each 50M mESCs before proceeding.

mESC pellets were thawed and resuspended in 750µL of FA-lysis buffer (50mM HEPES, 150mM NaCl, 2mM EDTA, 0.5mM EGTA, 0.5% IGEPAL CA-630, 0.1% sodium deoxycholate (Sigma-Aldrich D6750), 0.1% SDS). HEK293T cells were added and the combined cells incubated on ice for 10 minutes. Cells were sonicated on a Covaris E220e ultrasonicator (settings: peak incident power 140W, duty factor 5%, cycles per burst 200, temperature 4°C, run time 12.5min) in 1mL milliTUBEs with AFA Fiber (Covaris 520135) to shear chromatin. While the sonicator was running, 40µL Protein A Dynabeads™ (ThermoFisher 10008D) were washed twice with TE buffer, blocked at 4°C in TE buffer with 1mg/mL BSA, 1mg/mL yeast tRNA (Thermo Scientific AM7119) for 1 hour, washed once in TE buffer and once in FA-lysis buffer, and resuspended in 40µL FA-lysis buffer (80µL bead slurry per IP total). After sonication, cells were centrifuged at 20,000*g* and 4°C for 10 minutes; chromatin was taken as the supernatant.

For each immunoprecipitation reaction, 300µg of sonicated chromatin was diluted to 1mL with FA-lysis buffer and pre-cleared with 40µL of blocked bead slurry for 1 hour at 4°C. Cleared chromatin was spun down to remove clearing beads. 100µL of chromatin was set aside at 4°C as input. 15µL of anti-RPB1 antibody (Cell Signaling Technology 14958) was added to each IP and the mixture incubated overnight at 4°C. The next day, 40µL of blocked bead slurry was added to each IP and the mixture incubated for a further 3 hours at 4°C.

After bead binding, tubes were centrifuged at 1000*g* and 4°C for 5 minutes and supernatant removed. Beads were washed at 4°C for 5 minutes a total of five times: once with FA-lysis buffer, once with FA-lysis buffer with 500mM NaCl instead of 150mM, once with DOC buffer (10mM Tris-HCl pH 8, 250mM LiCl (Sigma-Aldrich L9650), 2mM EDTA, 0.5% IGEPAL CA-630, 0.5% sodium deoxycholate), and twice with TE buffer. All buffers were supplemented with 1X Roche cOmplete™ protease inhibitor cocktail, 1mM AEBSF (Fisher Scientific AC328110010), and 10mM NaF (Sigma-Aldrich 201154), and beads were centrifuged at 1000*g* and 4°C after each wash and supernatant removed. Washed beads were resuspended in 100µL of elution buffer (1% SDS, 0.1M $NaHCO_3$ (Sigma-Aldrich S6014)) and incubated at 30°C for 30 minutes with mixing at 1000rpm. Beads were centrifuged and the supernatant transferred to a new tube.

To reverse crosslinks, 4µL 5M NaCl and 2µL of 10mg/mL RNase A were added to each IP and input tube, and samples were incubated overnight at 65°C with shaking at 900rpm. The next day, 1µL of 20mg/mL proteinase K was added to each tube and samples incubated for 1 hour at 45°C. DNA was purified using a Zymo Clean and Concentrator kit following manufacturer's protocol. Purified DNA was post-sonicated on a Covaris E220e ultrasonicator (settings: peak incident power 105W, duty factor 5%, cycles per burst 200, temperature 4°C, run time 50s) in 130µL microTUBE AFA Fiber snap-cap tubes (Covaris 520045), then prepared for sequencing using a NEBNext Ultra II DNA library prep kit following manufacturer's protocol. Libraries were sequenced on an Illumina NovaSeq X.

**Acute perturbation of histone acetylation and calibrated native ChIP-seq**

To acutely perturb histone acetylation genome-wide, cells were treated with either 1µM trichostatin A (TSA, Sigma-Aldrich T8552) or 10µM A-485 (TargetMol T14073) for 4 hours. Perturbation of H3 acetylation was validated by Western blot[17] and profiled by spike-in calibrated native ChIP-seq[27–29,31].

To extract histones from treated cells for Western blots, cells were grown on 10cm dishes, harvested with TrypLE, washed with PBS, and resuspended in 1mL RSB buffer (10mM Tris-HCl pH 7.5, 10mM NaCl, 3mM $MgCl_2$) with 20mM N-ethylmaleimide (NEM, Fisher Scientific AA4052614). Cells were centrifuged at 1500rpm for 5 minutes and supernatant removed. Pelleted cells were resuspended in 1mL RSB with 0.5% IGEPAL CA-630 and 20mM NEM and incubated on ice for 10 minutes. Cells were centrifuged at 500*g* and 4°C for 5 minutes, then resuspended in 500µL of water with 5mM $MgCl_2$ and 20mM NEM. 500µL of 0.8M HCl (made from 1N HCl, Sigma-Aldrich H9892) was added and the mixture incubated on ice for 20 minutes, then centrifuged at 13,000rpm and 4°C for 20 minutes. The supernatant was transferred to a new tube. Trichloroacetic acid (TCA, Sigma-Aldrich T0699) was added to a final concentration of 25% v/v and the mixture incubated on ice for 30 minutes, then centrifuged at 13,000rpm and 4°C for 15 minutes. The supernatant was discarded. Pellets were washed twice in 1mL of ice-cold acetone (ThermoFisher 022928) and centrifuged at 13,000rpm and 4°C for 15 minutes for each wash. Pellets were air-dried for 15 minutes, resuspended in 80µL of 1X Laemmli loading buffer, and boiled at 90°C for 5 minutes. Western blotting was performed as described above, with the exception that fluorescent secondary antibodies were used and thus blots were not treated with Pierce™ ECL Western Blotting Substrate. Primary antibodies used: anti-histone H3 (Cell Signaling Technology 3638), anti-H3K27Ac (Abcam ab4729). Secondary antibodies used: IRDye 800CW anti-mouse IgG (LICORBio 926-32212), IRDye 680RD anti-rabbit IgG (LICORBio 926-68073).

Spike-in calibrated[27–29] native ChIP-seq of histone modifications was performed as previously described[31] with minor modifications. Three replicates were performed for each treatment. For each replicate, 50M mESCs and 20M *Drosophila* SG4 cells were combined and resuspended in 1mL RSB. 28mL of RSB with 0.1% IGEPAL CA-630 and 5mM sodium butyrate (Sigma-Aldrich 303410) and tubes inverted 10 times and centrifuged at 1500*g* and 4°C for 5 minutes. Cells were resuspended in 5mL RSB with 0.25M sucrose (Fisher Scientific BP220-1), 3mM $CaCl_2$, and 5mM sodium butyrate and centrifuged again as previous. Cells were then resuspended in 1mL RSB with 0.25M sucrose, 3mM $CaCl_2$, 5mM sodium butyrate, and 1X Roche cOmplete™ protease inhibitor cocktail. 150U of MNase was added to each sample, and the mixture incubated at 37°C for 5 minutes with shaking at 1000rpm. After 5 minutes, MNase digestion was immediately quenched with 4µL of 1M EDTA and the samples centrifuged at 5000rpm and 4°C for 5 minutes. The supernatant was removed as the S1 fraction and set aside. Pellets were resuspended in 300µL of nucleosome release buffer (10mM Tris-HCl pH 7.5, 10mM NaCl, 0.2mM EDTA) with 1X cOmplete™ protease inhibitor cocktail, 10mM NEM, and 5mM sodium butyrate and incubated at 4°C for 1 hour with rotation. Samples were passed 5 times through a 27G needle, then centrifuged at 5000rpm and 4°C for 5 minutes. 300µL of supernatant was removed as the S2 fraction and combined with the S1 fraction.

For each IP reaction, 100µL of combined chromatin was added to 900µL of native ChIP incubation buffer (70mM NaCl, 10mM Tris-HCl pH 7.5, 2mM $MgCl_2$, 2mM EDTA, 0.1% IGEPAL CA-630); an identical tube was made and set aside as input. 5µL of antibody (anti-H3K27Ac (Abcam ab4729) or anti-histone H3 (Cell Signaling Technology 3638)) was added to each IP and the tubes incubated at 4°C overnight. The next day, 20µL of pre-blocked Protein A beads (see above) were added and tubes incubated at 4°C for 1 hour. Beads were centrifuged at 1000*g* and 4°C for 1 minute and supernatant discarded. Beads were washed 4 times with 1mL of native ChIP wash buffer (20mM Tris pH 7.5, 2mM EDTA, 125mM NaCl, 0.1% IGEPAL CA-630) and once with 1mL TE buffer. Beads were resuspended in 100µL elution buffer. 100µL of each input sample was mixed with 20µL 10% SDS, 20µL 1M $NaHCO_3$, and 60µL of water and processed alongside the beads. All samples were incubated at room temperature for 30 minutes with shaking at 1000rpm, then centrifuged at room temperature and 1000*g* for 1 minute. DNA was purified from supernatants with a Zymo Clean and Concentrator kit following manufacturer's protocol. Libraries were prepared with a NEB Ultra II DNA library prep kit, individually indexed with NEBNext Unique UMI Dual-Index adaptors, and sequenced on an Illumina NovaSeq X.

**ChIP-seq data analysis**

Raw paired-end reads in .fastq format were aligned to a combined genome (mouse + spike-in) using the --no-mixed and --no-discordant options of bowtie2[32]. Duplicate reads were removed using sambamba markdup[33]. Aligned reads were separated into .bam files for mouse and spike-in genomes using samtools[34] and grep, and the output files were indexed with sambamba. Read counts were normalized with sambamba such that the final spike-in read counts were the same across samples[27–29]. BAM files were converted to BigWig files using a custom script employing MACS3[35] and the UCSC wigToBigWig utility. ChIP-seq profiles of the *Fbn2* locus were plotted using CoolBox[25]. Gene-body pileups of ChIP-seq signal were plotted using deepTools[36].

**ATAC-seq**

ATAC-seq was performed according to the protocol of Buenrostro *et al.*[37,38] with modifications. Briefly, 50,000 P2B2 cells were harvested with TrypLE, washed with PBS, and resuspended in 50µL of lysis buffer (10mM Tris-HCl, 10mM NaCl, 3mM $MgCl_2$, 0.1% IGEPAL CA-630, 0.1% Tween-20, 0.01% digitonin (Promega G9441)). Cells were incubated on ice for 3 minutes, then 1mL of wash buffer (10mM Tris-HCl, 10mM NaCl, 3mM $MgCl_2$, 0.1% Tween-20) was added to each tube and the mixture centrifuged at 500*g* and 4°C for 10 minutes. The supernatant was discarded. Nuclei were resuspended in 47.5µL of 1X TD Buffer (Illumina 20034198) with 0.1% Tween-20, 0.01% digitonin, and 2.5µL Tn5 transposase (Tagment DNA Enzyme 1, Illumina 20034198) and incubated at 37°C for 30 minutes with shaking at 1000rpm. DNA was purified using a Zymo DNA Clean and Concentrator kit following manufacturer's protocol.

After purification, DNA was PCR amplified for 10 cycles. Libraries were prepared with a NEB Ultra II DNA library prep kit, individually indexed with NEBNext Unique UMI Dual-Index adaptors, and sequenced on an Illumina NovaSeq X.

**ATAC-seq data analysis**

Raw paired-end reads in .fastq format were aligned to the mm39 genome using the --no-mixed and --no-discordant options of bowtie2[32]. Duplicate reads were removed using sambamba markdup[33]. BAM files were converted to BigWig files using MACS[35].

**Live-cell movie acquisition**

For live-cell imaging, 300,000-400,000 P2B2 cells were seeded in each well of an 8-well glass bottom chamber slide (no. 1.5H, Thermo Scientific 155409) that had been pre-coated with 0.1% gelatin for at least 24 hours two days prior to imaging. One day prior to imaging, medium was replaced with Dulbecco's Modified Eagle Medium (DMEM) without phenol red (Thermo Scientific 31053036) supplemented with 15% fetal bovine serum, 1000U/mL LIF, 1X GlutaMAX supplement, 1X MEM Non-Essential Amino Acid Solution, 0.1mM β-mercaptoethanol, 100μg/mL penicillin-streptomycin, and 2i: 1μM MEK inhibitor and 3μM GSK inhibitor. Cells were stained overnight with 50nM Halo-JFX650 dye; dye was not washed out before imaging. 3 hours before imaging, medium was supplemented with 1:100 ProLong Live Antifade Reagent (Thermo Scientific P36975). Transcriptional activation of *Fbn2*, RNAPII inhibition, and acute depletion of CTCF/RAD21 were performed as described above.

Live-cell imaging was conducted on a Zeiss Lattice Lightsheet 7 microscope fitted with a 3.3x/0.4NA objective for excitation and a 44.83x/1.0NA objective for detection, using a 20x850 light sheet. Zen 3.12 software was used to control incubation and acquisition. During imaging, cells were incubated at 5.5% $CO_2$, 37°C, and 85% relative humidity. Movies were taken at 30s intervals for a total of 3 hours (360 frames). Samples were loaded on the meniscus lens with water immersion, which was refreshed before beginning each movie. 500 slices were imaged per movie, with a step size between slices of 300nm. Three lasers were used, with a 15ms exposure time and the following power values: 488nm 4.0%, 561nm 7.5%, 640nm 10.0%. Data were acquired on two Hamamatsu ORCA-Fusion sCMOS cameras. The 488nm (corresponding to the R1 CTCF site) and 640nm (corresponding to MS2 on nascent transcripts) lasers were fired simultaneously, with the emissions split by an SBS LP 640 dichroic mirror. The MS2 signal was directed through a BP 570-620/LP 655 emission filter to camera 1, while the R1 signal was directed through a BP 495-550/BP 570-620 emission filter to camera 2. The 561nm laser (corresponding to the L1 CTCF site) was fired subsequently and emission captured on camera 2. At each time step, an image was acquired, then the microscope was allowed to auto-focus, using a custom macro.

To measure and correct for chromatic aberrations, a sample of 0.1μm TetraSpeck microspheres (ThermoFisher T2729) on mESCs was prepared and imaged frequently. 100,000 mESCs were plated in one well of an 8-well chamber slide and cultured as above. The next day, the vial of TetraSpeck beads was sonicated with a VEVOR Ultrasonic Cleaner for 5 minutes, and bead suspension was added to culture medium at a dilution of 1:100. The diluted bead suspension was sonicated for a further 10 minutes, then used to replace medium in the chamber slide. The following day, medium was replaced with 4% formaldehyde in PBS (diluted from 16% formaldehyde) and cells incubated for 15 minutes. Fixed cells were washed 3 times with PBS for 5 minutes per wash. After the final wash, 400μL PBS was added to cells. PBS was periodically replenished to prevent the sample from drying out. To record chromatic offset, we took 500-slice, 10-frame movies with 15ms exposure and 100% power for all three lasers. 4-5 such movies were taken each week of imaging.

**Live-cell movie processing**

Live-cell movies were processed using Fyrtarn[13] to extract 2-locus 3D distance and MS2 intensity trajectories as previously described[13]. Briefly, movies were deskewed and Richardson-Lucy deconvolution with total variation regularization applied. Point-spread function (PSF) kernels were computed from bead movies to enable deconvolution. The final pixel size of deskewed, deconvolved movies was 145x145x145nm. After deconvolution, nuclei were segmented from the Anchor3::OR3-2x-mStayGold-NLS signal via watershed-like erosion/dilation using CuPy[39]. Nuclear centroids were linked with TrackPy[40] with a search radius of 5.8μm and filtered for a minimum track length of 60 frames. CTCF site dots were detected using scale-space maxima detection. To minimize false positive detections at the nuclear edge, localizations falling within 1 pixel of the edge of the nucleus were removed. Localizations were subsequently refined with the FracShift algorithm[41] for 30 iterations. CTCF site detections were linked separately per-channel between frames, then joined between channels. Detections were linked between frames using TrackPy[40] with search radius 1450nm, memory of 5 frames, and minimum track length 15 frames. A Hungarian Algorithm[42]-like approach was used to pair localizations between channels.

To estimate chromatic aberrations, bead localizations were estimated from maxima of a Laplacian of Gaussian filter on bead movies and refined using FracShift. Localizations were paired with a search radius of 3 pixels and offset vectors recorded. To account for registration drifting over time, we fit a second-order polynomial correction function for each imaging day using bead localizations from a 15-day moving window centered on that day. Ridge regression with L2 weight penalty of $\lambda$=0.001 was performed to reduce overfitting. The correction functions were applied to detected CTCF site localizations before proceeding.

To estimate a location for MS2 intensity measurement, the position of the L1 CTCF site was estimated in the 640nm channel by combining cross-correlation-based image registration with chromatic aberration estimation as previously described. The dot detection process used for CTCF site dots was then applied to the 640nm channel. If a dot was detected within 5 pixels of the L1 CTCF site, MS2 intensity measurement was localized at that location; otherwise, the location of the L1 CTCF site was used. MS2 background signal was estimated by fitting a second-order polynomial with ridge regression (L2 weight penalty $\lambda$=0.001) and the background signal subtracted from MS2 intensity measurements. Photobleaching curves for the 640nm channel were computed from individual nuclei, normalized to the maximum intensity of each nucleus, and an ensemble photobleaching curve computed from the normalized curves for each movie.

Cross-channel dot linking and subsequent analyses can be confounded by DNA replication, which results in two highly proximal dots in a single channel that cannot be distinguished by the linkage algorithm and thus leads to faulty pairings[4,13]. To avoid this, replicated and false positive dots were removed by hand using the Fyrtarn quality control GUI. Trajectories with multiple neighboring detections in a single channel, large gaps of missing localizations in one channel, or poorly calibrated MS2 background signal were removed. Following manual QC, the squared Mahalanobis distances of frame-to-frame displacements were modeled using a chi-squared distribution with three degrees of freedom. Points exceeding a baseline significance level of $\alpha$=1e-7 with Bonferroni correction were masked (the second frame in the jump set to NaN) in the final data.

To infer periods of active transcription of *Fbn2* from MS2 intensity trajectories, we trained a 2-state hidden Markov model (HMM) with Gaussian emission distributions on MS2 data from the +ABA condition using the hmmlearn package[43]. To account for cell-to-cell variability in MS2 expression level, intensity trajectories were first z-score transformed using the mean value of MS2 trajectories from untreated cells as the mean and the per-nucleus standard deviation. Normalized intensities were then smoothed using a 5-frame rolling mean. Ten 2-state Gaussian HMMs were fit on the resulting data, and the model with the lowest Akaike information criterion (AIC) was selected as optimal. The model state with the higher mean was designated as "ON" and the other state as "OFF". Individual trajectories were labeled with transcription states as predicted by the HMM.

##### 3D distance calculation and Bayesian MSD fitting

3D distances between the anchors of the *Fbn2* TAD were computed as the Euclidean distance between paired localizations in the 488nm (R1) and 561nm (L1) channels of live-cell movies. Mean-squared displacement (MSD) analysis was performed on 3D distance trajectories using the two-locus heuristic fitting method implemented in the bayesmsd package[4]. The UNT and +ABA conditions were fit once with no fixed parameters to obtain an estimate of the localization error σ, then all conditions were refit with localization errors fixed to the mean values of the UNT and +ABA conditions to obtain values for anomalous exponent α and diffusion coefficient *D*.

##### Bayesian inference of looping probability and lifetime

To estimate the fully-looped probability and median loop lifetime of the *Fbn2* CTCF-CTCF loop, we applied Bayesian Inference of Looping Dynamics (BILD) as previously described[4,13]. Briefly, the algorithm defines a "looped" and "unlooped" state based on control experiments, then uses a hierarchical Bayesian model to assign those states to segments of experimental trajectories. We calibrated the model by fitting MSDs for ΔRAD21 and ΔCTCF 3D distance trajectories with α fixed to 0.5 (as BILD assumes a Rouse polymer, which has α=0.5). Parameters from the ΔCTCF fit were taken directly as the "unlooped" state. For the parameters of the "looped" state, we rescaled the parameters of the ΔRAD21 fit to match what would be expected from the ~7.5kb between the centers of the fluorescent DNA arrays when the *Fbn2* TAD CTCF sites were adjacent. These fitted parameters were used to calculate Rouse model parameters as previously described[4].

The calibrated BILD model was used to infer looping events in experimental trajectories. To prevent bias from varying trajectory length, we ran BILD on 100-frame subsamples of trajectories with starting frames offset by 50 frames (so frame 0-100, 50-150, 100-200, etc.). Only trajectories with at least 100 frames were analyzed. When two inferences for a single time point disagreed, the inference closer to the middle of its segment was preferred (e.g. a disagreement at frame 60 would prefer the inference from the 0-100 frame window over the 50-150 frame window). All BILD inferences were performed with evidence bias $\Delta E$ = 2, in accordance with previous work[4,13].

Looping probability was calculated for each condition as the fraction of timepoints where BILD predicted a loop, corrected as previously described[4] for false positives with $\Delta E$ = 2. Confidence intervals were estimated by track-based bootstrapping[4]. Loop lifetimes were estimated using the Kaplan-Meier survival estimator to account for censoring at the starts and ends of observed trajectories as in previous work[4].

##### Polymer simulations

Polymer simulations were performed using a 1D to 3D approach akin to previous work[4,13,44,45]. The code used here draws heavily from that employed by Goel *et al*[44]. First, cohesin and RNAPII positions were defined by a 1D fixed-timestep Monte Carlo simulation with 20 repeats of 2004 lattice sites each representing $\sigma$ = 500bp of chromatin. CTCF-binding sites (CBSs) were marked at single monomers in locations corresponding to the *Fbn2* locus (left-pointing: 132, 1166; right-pointing: 155, 1169, 1400, 1433, 1596). Upon encountering a CBS, a cohesin motor would either stall with probability $s$ = 0.8 or bypass the CBS with probability 0.2, in accordance with previous work[4]. Once a motor stalled, that motor could not move further until it dissociated from chromatin and bound again elsewhere. The other arm of that cohesin could continue moving. CBSs in these simulations were directional and would only stall cohesin if they were convergent with the direction of extrusion.

All 1D simulations were performed with a fixed number of cohesins determined by the ratio of chromosome length $G$ and cohesin separation $d$ = 480 monomers (240 kb), as reported by Gabriele *et al.*[4]. At each timestep, each cohesin motor subunit had a chance to move defined as $v_{LEF}$ and a chance to dissociate from chromatin defined as $1/L$, where $L$ is cohesin lifetime. $L$ and $v_{LEF}$ were determined by a parameter sweep to maximize correspondence to the experimentally observed looping probability of the *Fbn2* TAD. Optimal values used for all simulations were $L$ = 2150 timesteps and $v_{LEF}$ = 0.125.

Genes were marked at the following monomer locations: starts = 622, 626, 1168, 1521, 1875; ends = 219, 650, 1170, 1561, 2003. For single-model simulations, promoters (start monomers) were constitutively active, such that at each timestep an RNAPII could associate with the lattice with probability $k_{init}$ = 0.0167 and process until reaching the corresponding end site, at which point it would stall until dissociating. For simulations with moving barrier activity, cohesins would have a probability to bypass an RNAPII of $s_{pol}$, which was defined for each simulation. If a cohesin subunit failed to bypass RNAPII, it would remain stationary until the RNAPII moved, at which point it could again attempt to bypass the next time that cohesin subunit moved. As with CTCF, the other cohesin subunit could continue moving as normal. Each RNAPII had a chance to move once per timestep of $v_{pol}$ = 0.016667. When RNAPII moved, it would push any cohesin subunits present in the direction of motion one monomer backwards.

Next, the 1D simulation was coupled to a 3D polymer model. Molecular dynamics simulations were performed using Polychrom[46], which wraps the OpenMM simulation toolkit[47]. Each cohesin was modeled in the 3D simulation as a harmonic bond between two monomers, and their locations were dynamically updated based on the positions of cohesins in the 1D simulation. The 1D simulation was run for 80,000 timesteps to equilibrate LEF positions before coupling to the 3D simulation. As in the 1D simulation, the system was modeled as a chromosome of $G$ = 40,080 sites, containing 20 repeats of a 2004-monomer locus corresponding to the *Fbn2* locus. Each monomer was assigned one of three types (A, B, or C), and could interact with other monomers through homotypic ($\varepsilon_A$, $\varepsilon_B$, and $\varepsilon_C$) and heterotypic ($\varepsilon_{BC}$, B-C interactions only) attractive interaction. Interaction potentials between monomers were given by a smooth square-well potential, as in previous work[44]. Values of homotypic and heterotypic attractions were modulated based on the model being tested (Table S4). C-type monomers were placed in 3-monomer groups at gene start sites. For the local decompaction model, a baseline simulation was performed with B-type monomers everywhere except C-type monomers and monomers upstream of the L1 CTCF site (monomer 155), then monomers 155-622 were switched to A-type and the simulation repeated. For the local self-affinity model, monomers upstream of the *Fbn2* start site (0-622) were set to A-type, with the rest of the non-C-type monomer set to B-type. For the microcompartment and moving barrier model, all non-C monomers were set to B-type.

Simulations were confined to a spherical region with prescribed chromatin density $\rho$ = 0.2. Simulations were evolved using the OpenMM fixed timestep Langevin integrator with timestep = 40 and collision_rate = 0.01, with 450 polymer time steps between each loop extrusion update step. 1D and 3D simulation times and distances were calibrated by measuring the 2-point root mean squared displacement (RMSD) and Rouse prefactor $\Gamma$ of the simulated *Fbn2* TAD and comparing the values to those determined experimentally in previous work[4]. This resulted in a monomer size of $a$ = 15nm and a polymer timestep of 1.1ms; 450 polymer steps per loop extrusion step were conducted such that each loop extrusion step corresponded to 0.5s. 1,000,000 loop extrusion steps were conducted for each simulated condition, with conformations saved every 100 steps for a total of 10,000 saved conformations (200,000 individual locus conformations) for each condition.

Polymer simulations of the combined mechanistic model were conducted largely as above. In these simulations, transcriptional bursting was implemented such that RNAPII could only associate with chromatin when the system was in an ON state. At each timestep, the system could move from OFF to ON with probability $k_{on}$ or from ON to OFF with probability $k_{off}$ = 1.667e-3. We conducted a parameter sweep to calibrate $k_{on}$, $s_{pol}$, and $\varepsilon_B$ to best match experimental observations (Fig. S10, Table S4). To simulate only local decompaction, $s_{pol}$ was set to 1. To simulate RAD21 depletion, $v_{LEF}$ was set to 1e-11 and *d* set to 4e13. To simulate only moving barrier activity, monomers between the L1 CTCF site and the *Fbn2* promoter (monomers 155-622) were not switched from B-type to A-type.

Contact maps for simulated polymer configurations were computed and plotted using polychrom[46]. Simulated contact maps were balanced by applying the Sinkhorn-Knopp algorithm[48]. Insulation scores were computed from simulated contact maps as above. MSDs for simulated trajectories were calculated with the noctiluca package[4].

### METHODS REFERENCES

39. Okuta, R., Unno, Y., Nishino, D., Hido, S. & Loomis, C. CuPy: A NumPy-Compatible Library for NVIDIA GPU Calculations.
40. Allan, D. B., Caswell, T., Keim, N. C., van der Wel, C. M. & Verweij, R. W. soft-matter/trackpy: v0.7. https://doi.org/10.5281/zenodo.16089574 (2025) doi:10.5281/zenodo.16089574.
41. Gao, Y. & Kilfoil, M. L. Accurate detection and complete tracking of large populations of features in three dimensions. *Opt. Express* **17**, 4685–4704 (2009).
42. Kuhn, H. W. The Hungarian method for the assignment problem†. *Nav. Res. Logist. Q.* **2**, 83–97 (1955).
43. Pedregosa, F. *et al.* Scikit-learn: Machine Learning in Python. *J. Mach. Learn. Res.* **12**, 2825–2830 (2011).
44. Goel, V. Y. *et al.* Dynamics of microcompartment formation at the mitosis-to-G1 transition. *Nat. Struct. Mol. Biol.* **32**, 2614–2627 (2025).
45. Banigan, E. J. *et al.* Transcription shapes 3D chromatin organization by interacting with loop extrusion. *Proc. Natl. Acad. Sci.* **120**, e2210480120 (2023).
46. Imakaev, M., Goloborodko, A. & hbbrandao. mirnylab/polychrom: v0.1.0. https://doi.org/10.5281/zenodo.3579473 (2019) doi:10.5281/zenodo.3579473.
47. Eastman, P. & Pande, V. S. OpenMM: A Hardware Independent Framework for Molecular Simulations. *Comput. Sci. Eng.* **12**, 34–39 (2015).
48. Knopp, P. & Sinkhorn, R. Concerning nonnegative matrices and doubly stochastic matrices. *Pac. J. Math.* **21**, 343–348 (1967).

#### SUPPLEMENTARY FIGURES AND FIGURE LEGENDS

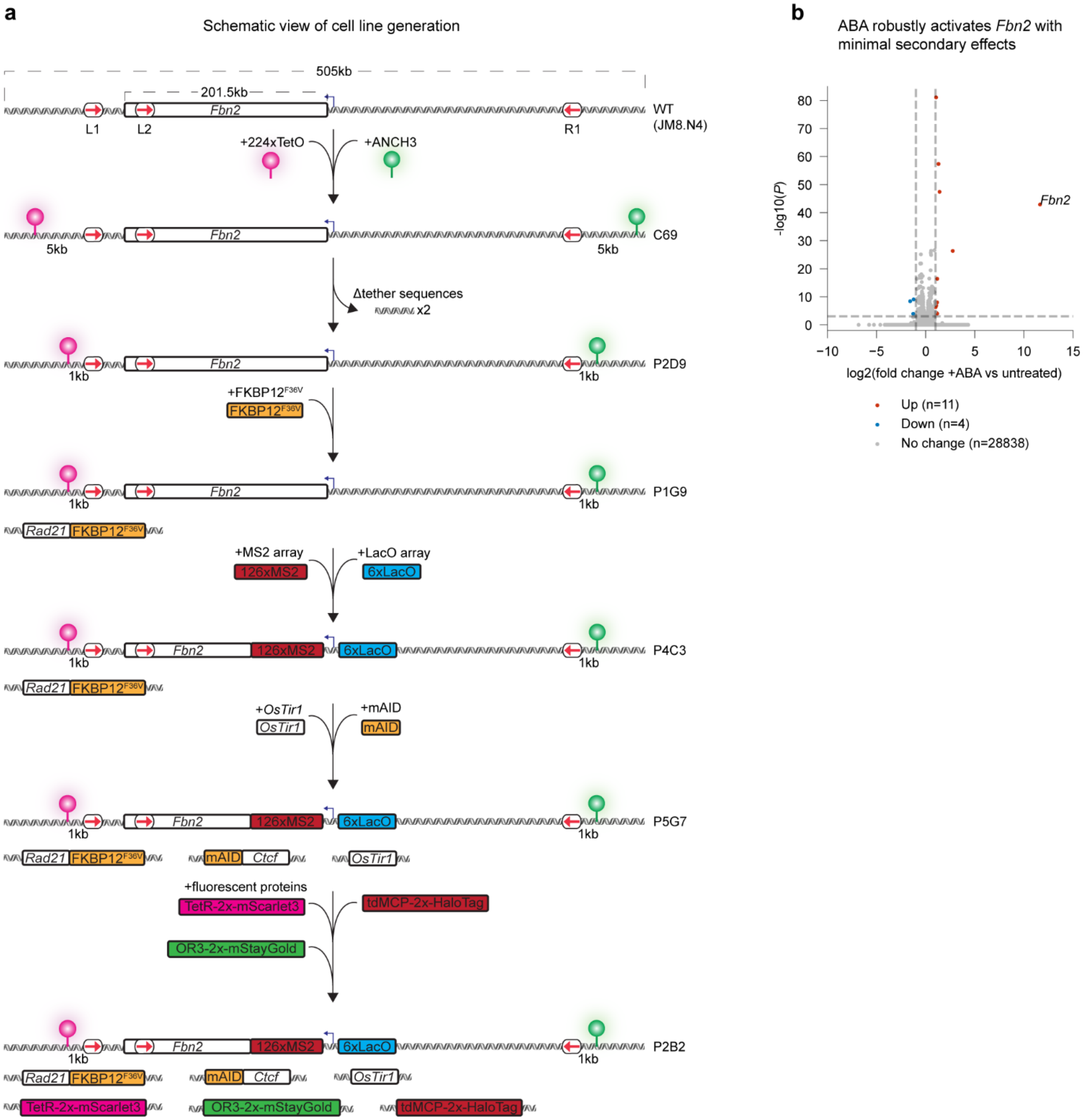


**Figure S1: Developing a system for investigating 3D genome impacts of transcription. a,** Schematic overview of genome editing process to create the P2B2 cell line used in this work. **b,** Differentially expressed genes in +ABA vs untreated cells, measured by RNA-seq. *Fbn2* is annotated.

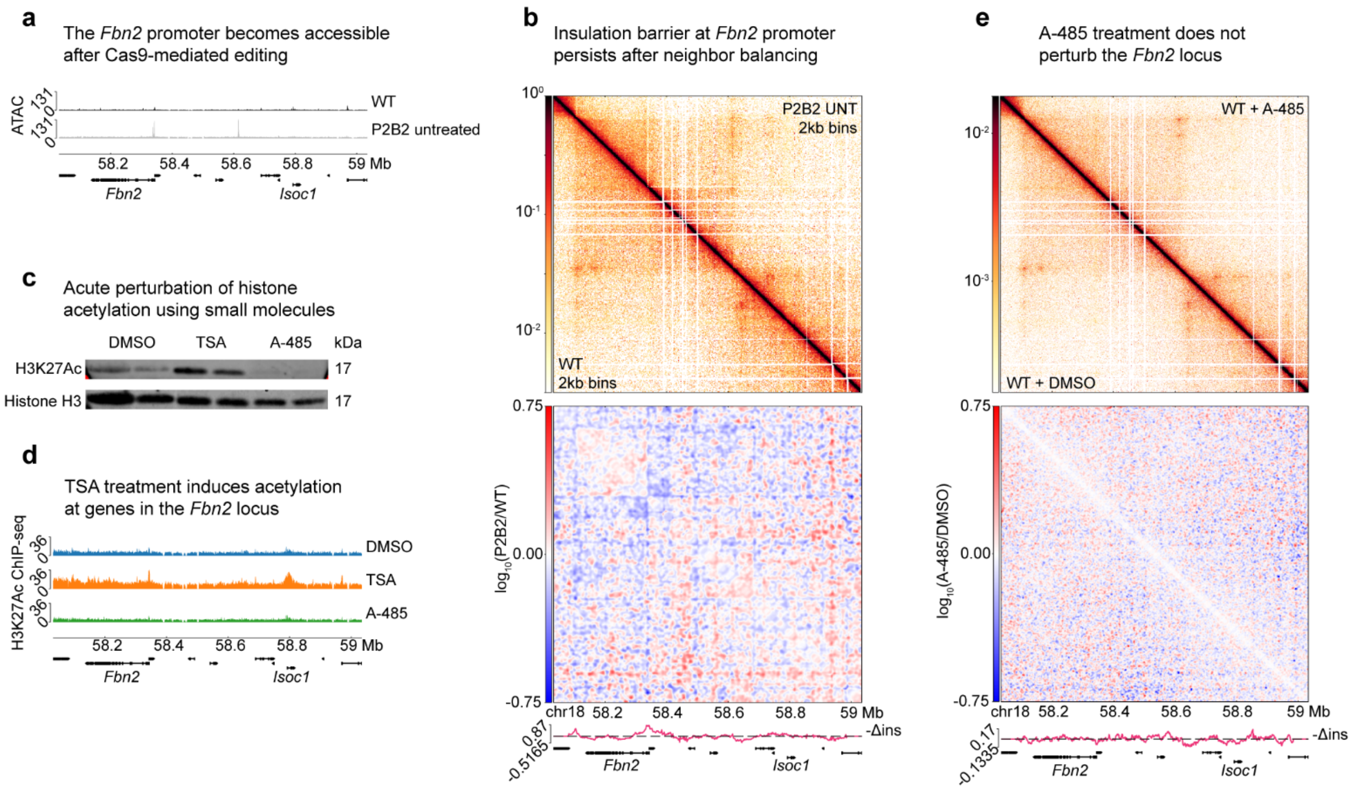


**Figure S2: Investigating genome editing-associated insulation at the *Fbn2* promoter. a,** ATAC-seq tracks across the *Fbn2* locus, before and after genome editing. **b,** Neighbor-balanced raw contact maps (top right: P2B2, bottom left: WT), differential contact map (P2B2/WT), and differential insulation score (WT - P2B2) of the *Fbn2* locus. **c,** Western blots showing acute perturbation of H3K27Ac by TSA and A-485, with histone H3 as loading control. **d,** H3K27Ac ChIP-seq tracks across the *Fbn2* locus for WT mESCs treated with DMSO, TSA, or A-485 for 4 hours. **e,** As **b**, with WT +A-485 cells in top right and WT +DMSO cells in bottom left.

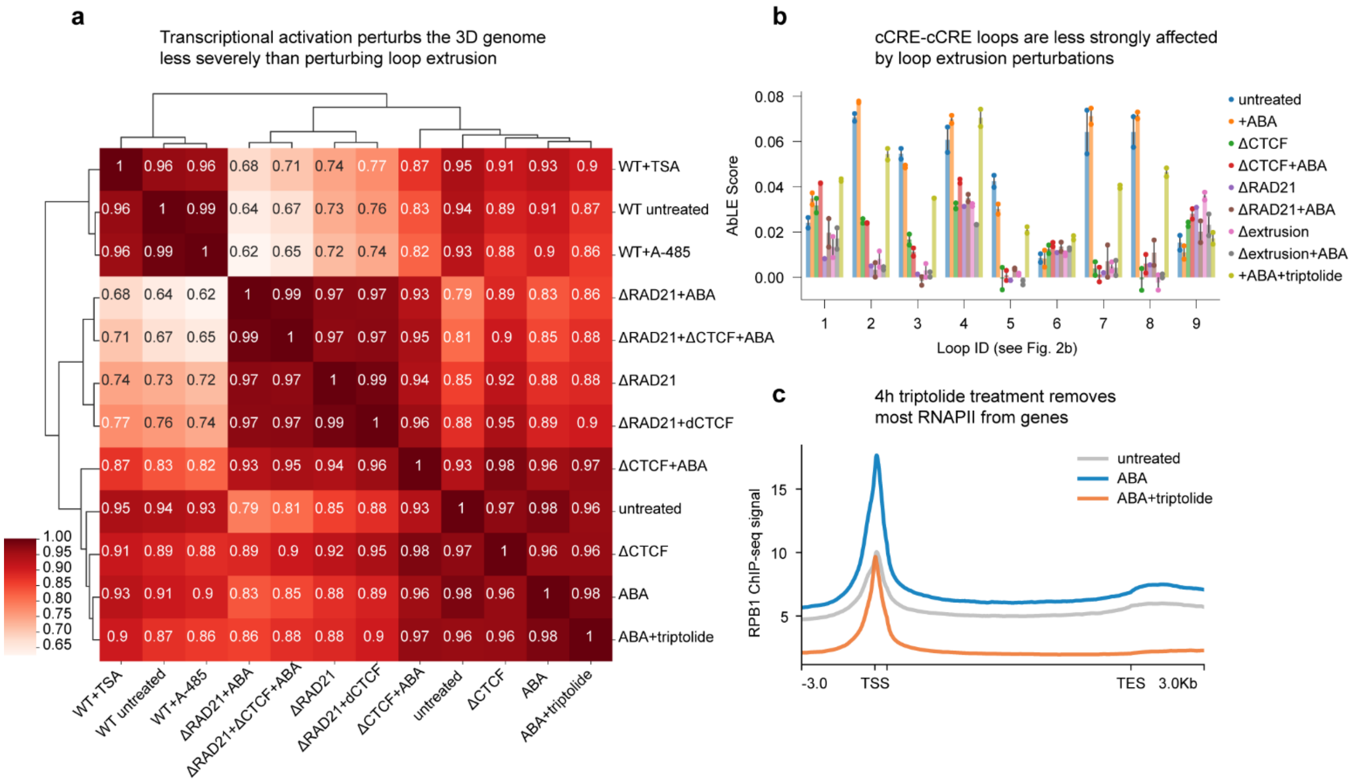


**Figure S3: Transcriptional activation affects contact patterns but not loops in the *Fbn2* locus. a,** Stratum-adjusted correlation coefficient (SCC) between RCMC contact maps of *Fbn2* locus across all conditions in this study. **b,** AbLE scores for key loops in the *Fbn2* locus in P2B2 cells across all perturbations. **c,** RNAPII ChIP-seq signals averaged across all gene bodies.

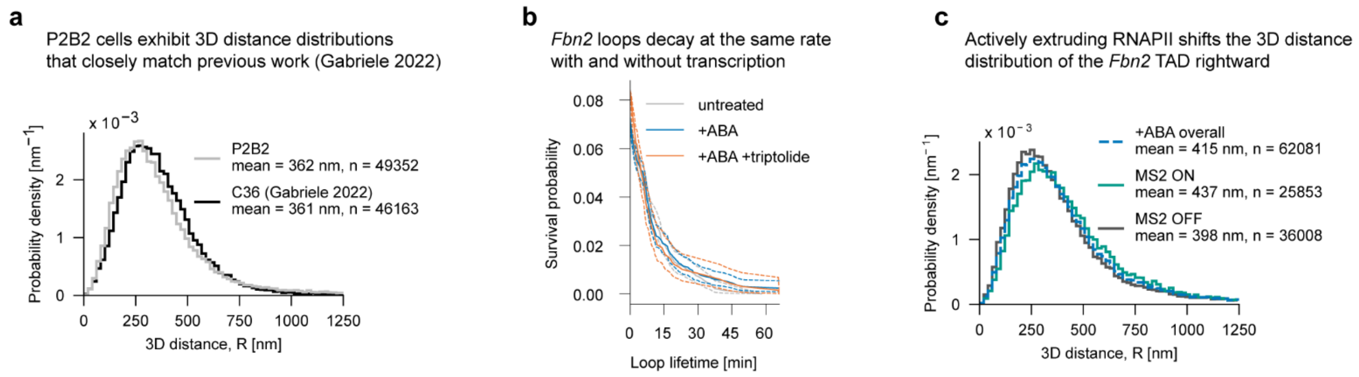


**Figure S4: Profiling the dynamic impact of transcriptional activation. a,** 2-point 3D distance distributions between the ends of the *Fbn2* TAD for untreated P2B2 cells and untreated C36 cells from Gabriele *et al*[4]. **b,** Kaplan-Meier survival curves for the fully looped state of the *Fbn2* TAD. **c,** 2-point 3D distance distributions of the *Fbn2* TAD, stratified by MS2 burst state.

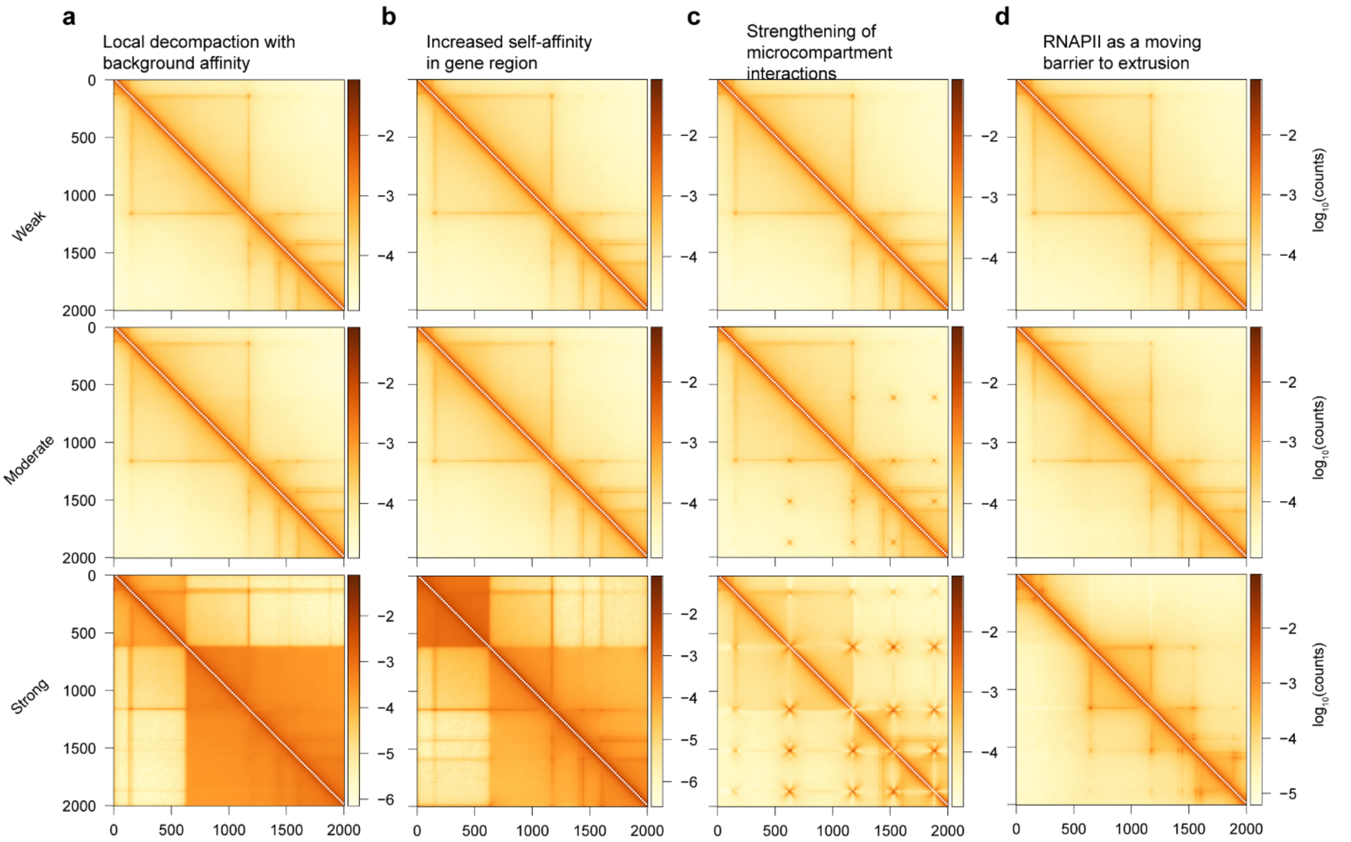


**Figure S5: Determining the 3D signatures of possible models via polymer simulations. a-d,** ICE-balanced contact maps of the simulated *Fbn2* TAD across multiple strengths of simulated perturbations.

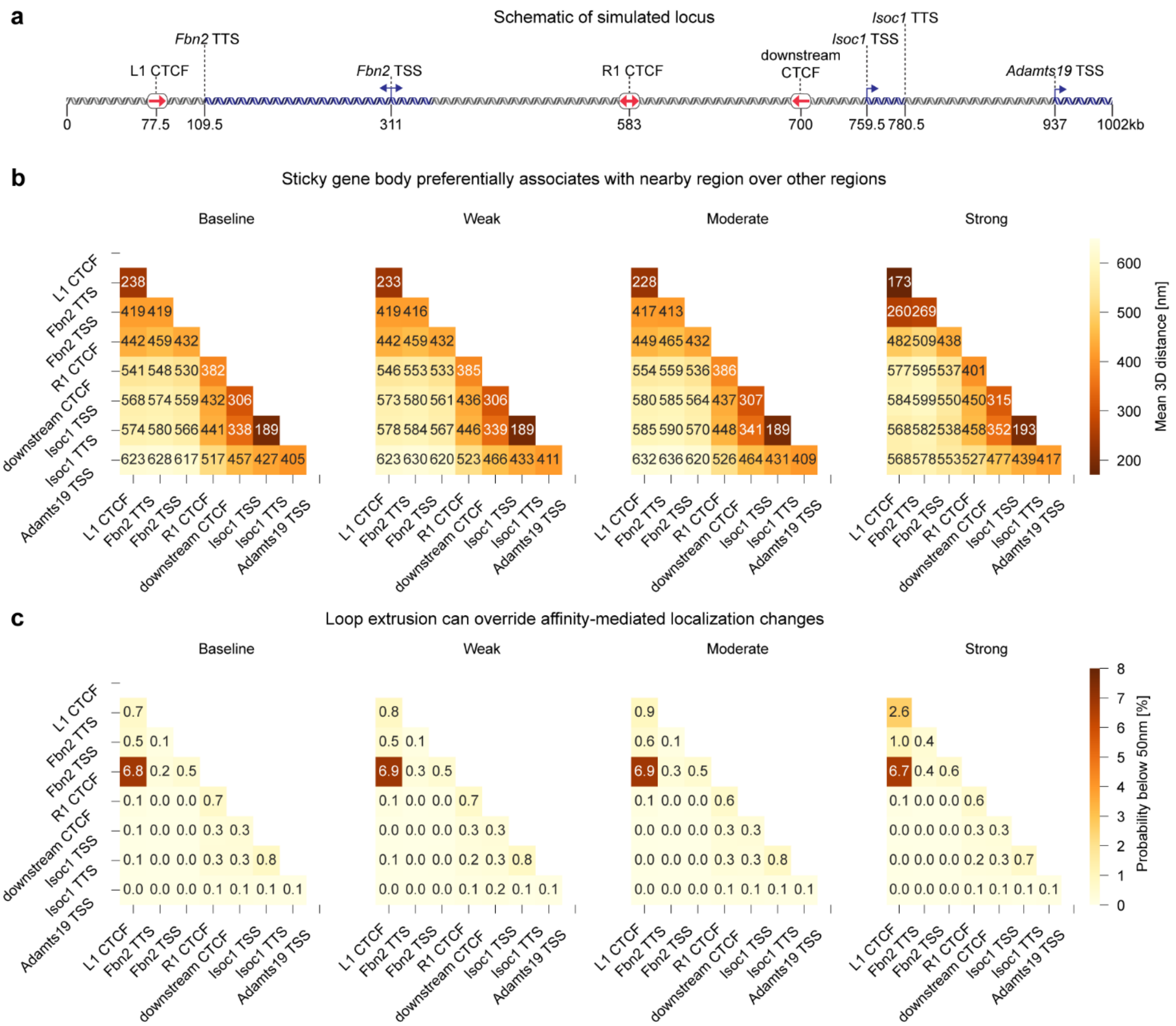


**Figure S6: Characterizing the sticky gene body model. a,** Schematic overview of the simulated *Fbn2* locus. **b,** Heatmap of mean 3D distances between key points in the simulated locus across the range of sticky gene body parameter strengths. **c,** Heatmap of probabilities that two points are within 50nm of each other for key points in the simulated locus across the range of sticky gene body parameter strengths.

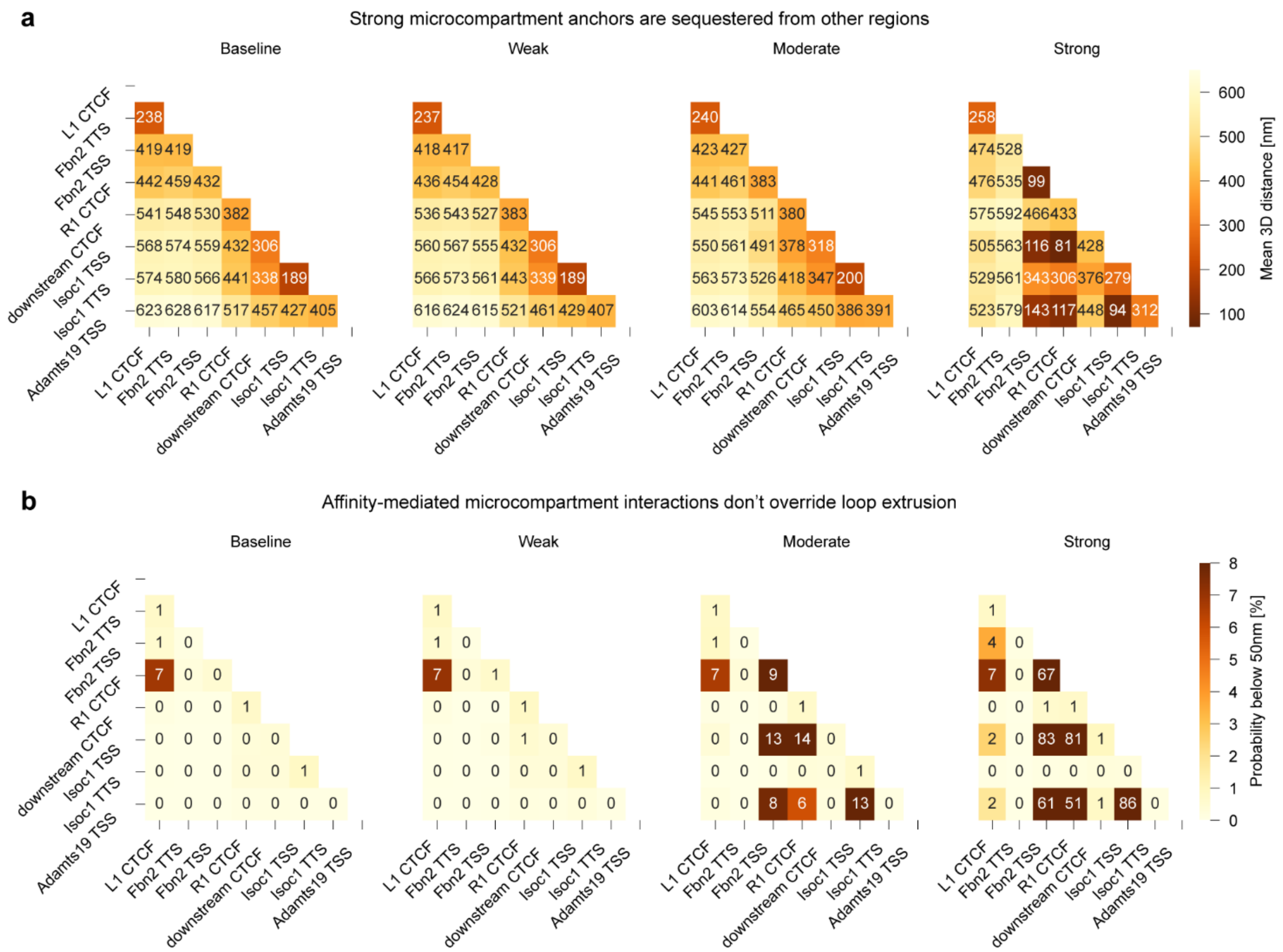


**Figure S7: Characterizing the strong microcompartment model. a,** Heatmap of mean 3D distances between key points in the simulated locus across the range of microcompartment parameter strengths. **b,** Heatmap of sub-50nm-distance probabilities for key points in the simulated locus across the range of microcompartment parameter strengths.

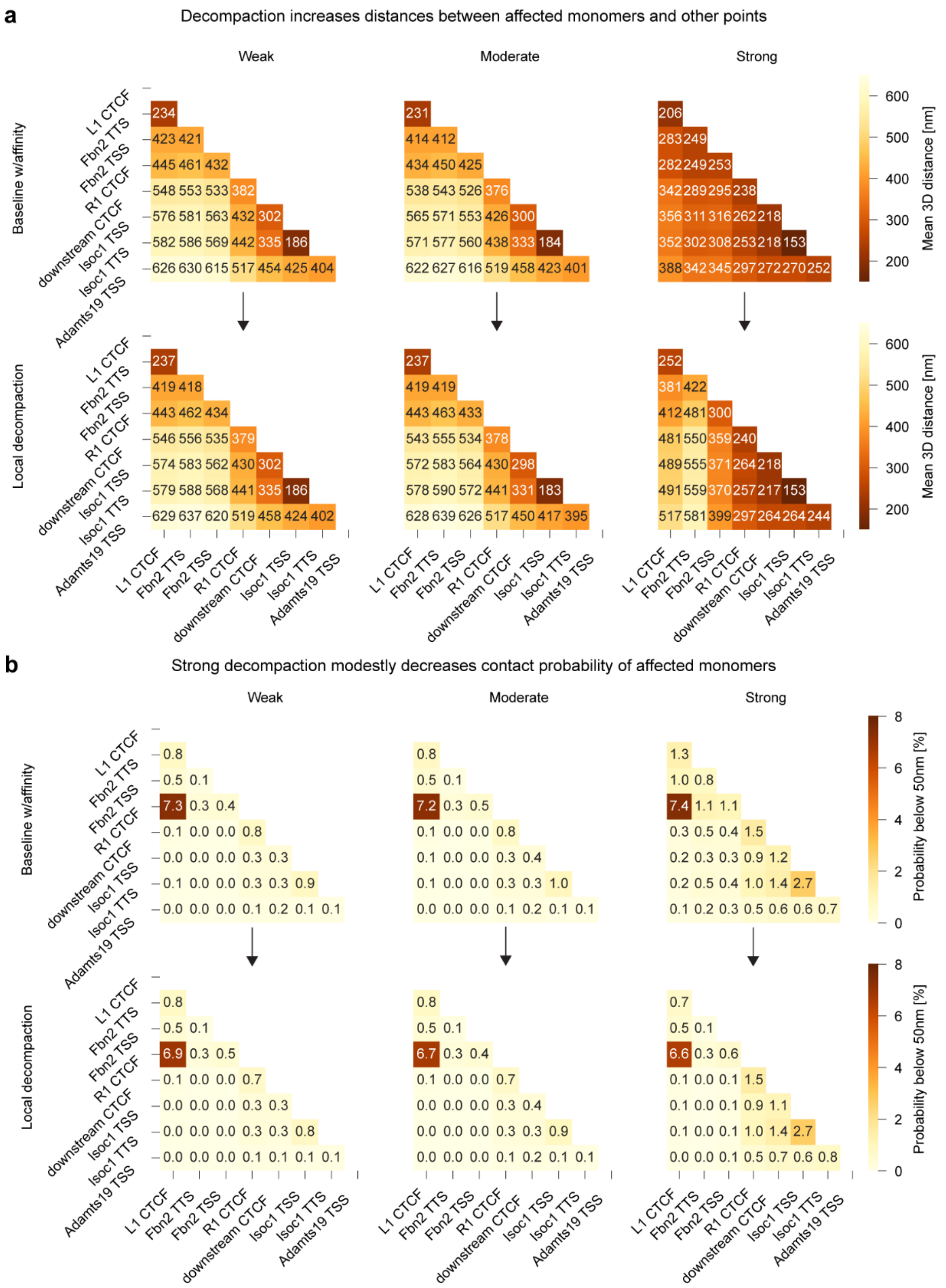


**Figure S8: Characterizing the decompaction model. a,** Heatmap of mean 3D distances between key points in the simulated locus across the range of local decompaction parameter strengths. Top row: baseline simulation with affinity at all B-type monomers. Bottom row: perturbed simulation with affinity removed from *Fbn2* region. **b,** Heatmap of sub-50nm-distance probabilities for key points in the simulated locus across the range of local decompaction parameter strengths. Top row: baseline simulation with affinity at all B-type monomers. Bottom row: perturbed simulation with affinity removed from *Fbn2* region.

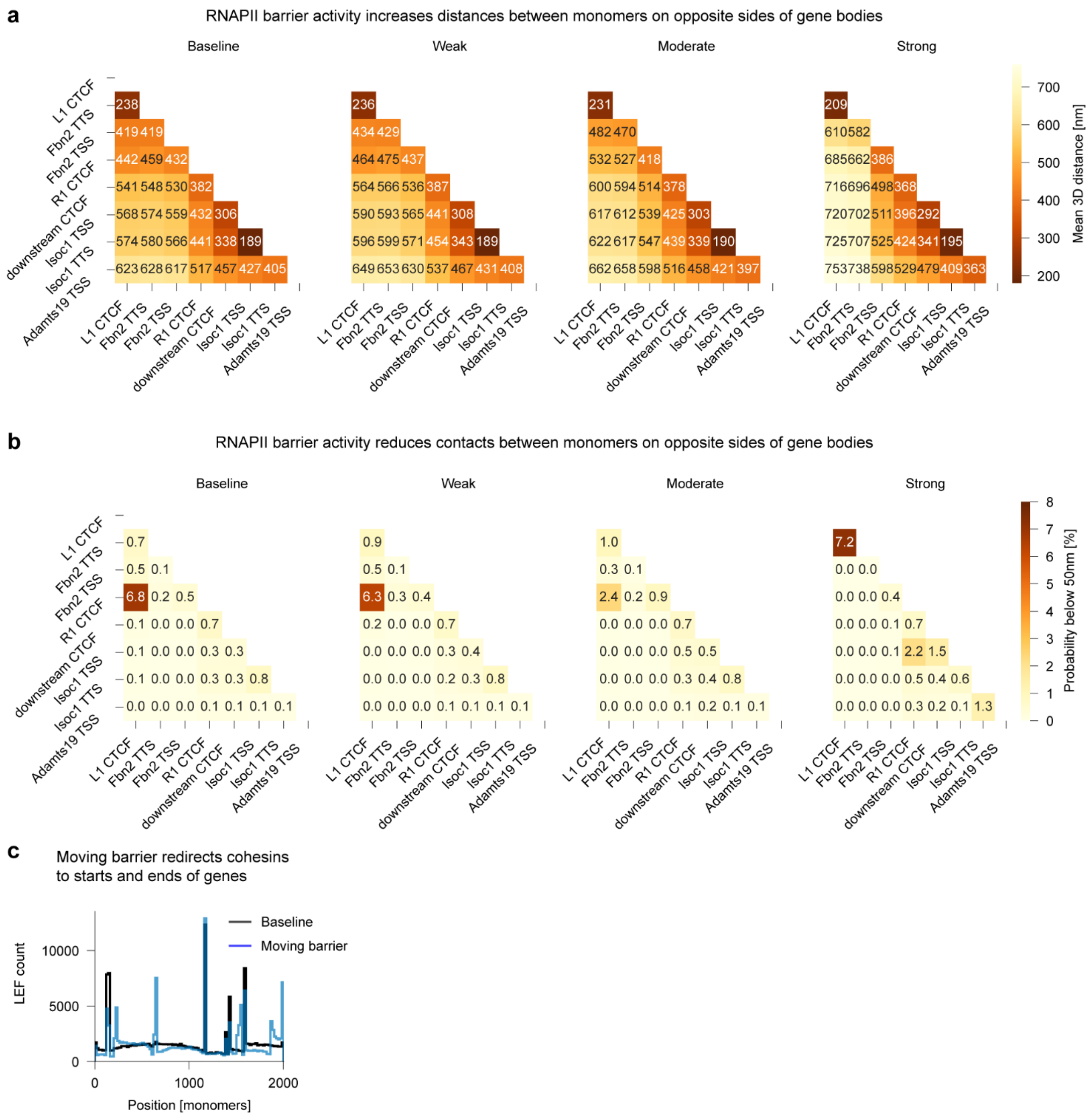


**Figure S9: Characterizing the moving barrier model. a,** Heatmap of mean 3D distances between key points in the simulated locus across the range of moving barrier parameter strengths. **b,** Heatmap of contact probabilities for key points in the simulated locus across the range of moving barrier parameter strengths. **c,** Pileup of LEF positions across all conformations of the simulated locus, for baseline and strong moving barrier conditions.

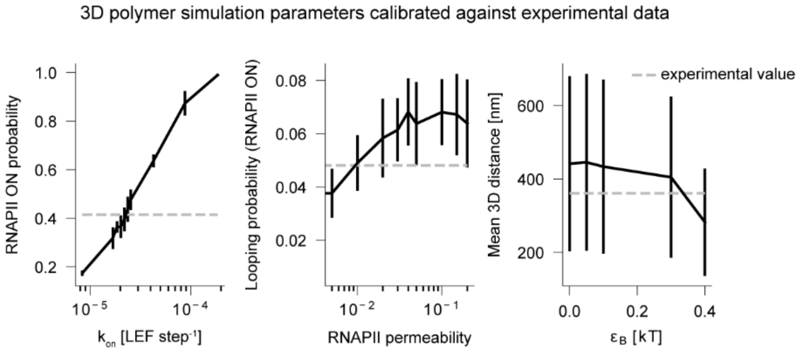


**Figure S10: Calibrating a combined mechanistic model.** Plots of RNAPII ON probability vs $k_{ON}$, looping probability during RNAPII presence vs RNAPII bypass probability, and mean 2-point 3D distance of the *Fbn2* TAD vs $\varepsilon_B$. Swept simulation parameters indicated with solid lines; experimental values shown by dashed lines. Error bars represent standard deviation.

#### SUPPLEMENTAL TABLES

**Table S1: Region-Capture Micro-C read counts**

| Condition | Replicate | Unique reads in capture region | Genome-wide equivalent unique reads |
|---|---|---|---|
| P2B2 untreated | 1 | 962,273 | 2,549,374,403 |
| | 2 | 436,912 | 1,157,522,105 |
| P2B2 +ABA | 1 | 1,571,103 | 4,162,363,250 |
| | 2 | 1,793,259 | 4,750,926,808 |
| P2B2 ΔCTCF | 1 | 749,255 | 1,985,020,382 |
| | 2 | 933,660 | 2,473,569,252 |
| P2B2 ΔCTCF+ABA | 1 | 2,904,868 | 7,695,940,884 |
| | 2 | 2,315,631 | 6,134,860,271 |
| P2B2 ΔRAD21 | 1 | 3,411,836 | 9,039,064,137 |
| | 2 | 5,444,489 | 14,424,223,575 |
| P2B2 ΔRAD21+ABA | 1 | 3,473,124 | 9,201,435,999 |
| | 2 | 4,308,517 | 11,414,663,981 |
| P2B2 ΔRAD21+ΔCTCF | 1 | 1,864,801 | 4,940,464,853 |
| | 2 | 1,836,354 | 4,865,099,490 |
| P2B2 +ABA+ΔRAD21+ ΔCTCF | 1 | 2,802,289 | 7,424,175,723 |
| | 2 | 2,654,286 | 7,032,067,600 |
| P2B2 +ABA+triptolide | 1 | 1,774,290 | 4,700,671,752 |
| | 2 | 2,314,982 | 6,133,140,859 |
| P2B2 +ABA+IPTG | 1 | 3,218,227 | 8,526,130,875 |
| | 2 | 2,469,329 | 6,542,056,302 |
| WT +DMSO | 1 | 377,976 | 1,001,381,457 |
| | 2 | 590,961 | 1,565,648,050 |
| WT +TSA | 1 | 750,837 | 1,989,211,615 |
| | 2 | 1,834,820 | 4,861,035,425 |
| WT +A-485 | 1 | 293,195 | 776,768,992 |
| | 2 | 748,034 | 1,981,785,556 |

**Table S2: Genome loop coordinates in the edited *Fbn2* locus**

| Loop ID | Start 1 | End 1 | Start 2 | End 2 |
|---|---|---|---|---|
| 1 | 58,101,069 | 58,106,069 | 58,338,957 | 58,343,957 |
| 2 | 58,105,046 | 58,110,046 | 58,612,352 | 58,617,352 |
| 3 | 58,165,713 | 58,170,713 | 58,612,352 | 58,617,352 |
| 4 | 58,339,643 | 58,344,643 | 58,608,346 | 58,613,346 |
| 5 | 58,104,052 | 58,109,052 | 58,726,639 | 58,731,639 |
| 6 | 58,336,661 | 58,341,661 | 58,787,276 | 58,792,276 |
| 7 | 58,633,853 | 58,638,853 | 58,726,639 | 58,731,639 |
| 8 | 58,633,853 | 58,638,853 | 58,737,543 | 58,742,543 |
| 9 | 58,612,014 | 58,617,014 | 58,788,270 | 58,793,270 |

**Table S3: Experimental MSD fit parameters**

| Condition | α | D [μm²*s⁻α] | 2<*R*²> [μm²] | σx [nm] | σy [nm] | σz [nm] |
|---|---|---|---|---|---|---|
| Untreated | 0.306 | 0.0163 | 0.36 | 38.2 | 30.7 | 54.6 |
| +ABA | 0.302 | 0.0217 | 0.49 | | | |
| +ABA+triptolide | 0.297 | 0.0276 | 0.55 | | | |
| ΔRAD21 | 0.312 | 0.0488 | 0.88 | | | |
| ΔCTCF | 0.322 | 0.0223 | 0.44 | | | |

**Table S4: Polymer simulation parameters**

| Condition | Strength | $\varepsilon_A$ [kT] | $\varepsilon_B$ [kT] | $\varepsilon_C$ [kT] | RNAPII bypass probability | $k_{ON}$ | $k_{OFF}$ |
|---|---|---|---|---|---|---|---|
| Local decompaction | weak | 0 | 0.1 | 0.1 | 1 | N/A | N/A |
| | moderate | 0 | 0.2 | 0.2 | 1 | N/A | N/A |
| | strong | 0 | 0.4 | 0.4 | 1 | N/A | N/A |
| Increased gene-body affinity | weak | 0.1 | 0 | 0 | 1 | N/A | N/A |
| | moderate | 0.2 | 0 | 0 | 1 | N/A | N/A |
| | strong | 0.4 | 0 | 0 | 1 | N/A | N/A |
| Strengthened microcompartment interactions | weak | 0 | 0 | 2 | 1 | N/A | N/A |
| | moderate | 0 | 0 | 3 | 1 | N/A | N/A |
| | strong | 0 | 0 | 4 | 1 | N/A | N/A |
| RNAPII moving barrier | weak | 0 | 0 | 0 | 0.1 | N/A | N/A |
| | moderate | 0 | 0 | 0 | 0.05 | N/A | N/A |
| | strong | 0 | 0 | 0 | 0 | N/A | N/A |
| combined | N/A | 0 | 0.33 | 0.33 | 0 | 2.195e-5 | 1.667e-3 |

**Table S5: List of plasmids used in this study**

| Name | Purpose | Corresponding sgRNA sequence(s) |
|---|---|---|
| pASH83 | Repair construct for inserting 224xTetO array upstream of L1 CTCF site of *Fbn2* TAD | GTTAACTGAGATCTATTCGC, TTAACTGAGATCTATTCGCA, GATCTATTCGCAGGGAACTA |
| pASH89 | Repair construct for inserting ANCH3 array downstream of R1 CTCF site of *Fbn2* TAD | TACCTTAGCACTGCCTCGTA, AAGTTTTGTGAGTCCTTACG, GTCCTTACGAGGCAGTGCTA |
| pASH360 | Cas9 expression vector with two sgRNA scaffolds | L1 tether deletion: TCCTGCACGACGCGAGCTGC and AGAGATCACCCCAATGGCAG; R1 tether deletion: GAAGAAGATTGTGAATGGCA and TCCTCTAGCAAAGAAAATGC |
| pASH247 | Repair construct for C-terminal tagging of *Rad21* with FKBP12$^{F36V}$ and V5 epitope tag | CCTCAGATAATATGGAACCG |
| pASH307 | Repair construct for N-terminal tagging of *Ctcf* with mini-AID tag and 3xFLAG epitope tag | TGAGGCGGTTGAAGCCATTG |
| pASH361 | Repair construct for inserting 126xMS2 array and 6xLacO array at the *Fbn2* promoter region | AAAGCTGCAGAAGTCACCAA and ACAGGTCCGACCCGCAGTAG |
| pASH362 | Repair construct for inserting PYL1-VPR and ABI-LacI constructs at the *Hipp11* locus[8] | AACACTAGTGCACTTATCCT |
| pASH424 | PiggyBac integration vector for expression of TetR-2xmScarlet3 construct | N/A |
| pASH380 | PiggyBac integration vector for expression of tdMCP-2xHaloTag construct | N/A |
| pASH133 | PiggyBac integration vector for expression of OsTIR1 and blasticidin resistance | N/A |
| pASH349 | PiggyBac integration vector for expression of OR3-2x-m(n1)oxStayGold construct | N/A |

**Table S6: List of oligonucleotides used in this study**

| Name | Sequence | Purpose |
|---|---|---|
| L1_geno_forward | GAGCACCTCATCGGCGATAG | Primer for screening the L1 tether deletion |
| L1_geno_reverse | TCCCTCAATCCCCTCCACAT | Primer for screening the L1 tether deletion |
| R1_geno_forward | TCAGCTTAGGGGTAGGGTTGT | Primer for screening the R1 tether deletion |
| R1_geno_reverse | GGCTCTGCTTTTGTCTTGGC | Primer for screening the R1 tether deletion |
| MS2_screening_F1 | CAGAAGGTGGAGTGCTGGAG | Primer for screening 126xMS2 and 6xLacO insertions at *Fbn2* promoter |
| MS2_screening_R1 | CCTAGCTGCCCCAAACGAG | Primer for screening 126xMS2 and 6xLacO insertions at *Fbn2* promoter |
| MS2_screening_F2 | TTGCGGACCCGTTACTACTG | Primer for screening 126xMS2 and 6xLacO insertions at *Fbn2* promoter |
| CTCF_screening_F | GGTGTCATTGTGTGGCCAAG | Primer for screening mAID tagging of *Ctcf* |
| CTCF_screening_R | GGCTGGTGGCTGATAGTCTG | Primer for screening mAID tagging of *Ctcf* |
| RAD21_screening_F1 | TTTCTGCACATGGGGATGGA | Primer for screening $FKBP12^{F36V}$ tagging of *Rad21* |
| RAD21_screening_R | TGAGGAGTCACGCCACTGTA | Primer for screening $FKBP12^{F36V}$ tagging of *Rad21* |
| RAD21_screening_F2 | GATCCGAGGCTGGGAAGAAG | Primer for screening $FKBP12^{F36V}$ tagging of *Rad21* |
| VPR_screening_F1 | TCTTTCCCTTGCCTCTGCTG | Primer for screening PYL1-VPR and ABI-LacI insertions at *Hipp11* locus |
| VPR_screening_R2 | ACTAGTCACCGCCACCATGGGTGG | Primer for screening PYL1-VPR and ABI-LacI insertions at *Hipp11* locus |
| VPR_screening_F2 | GCACCGATCGCTGCAGGTAATTCTACCGGGTAGG | Primer for screening PYL1-VPR and ABI-LacI insertions at *Hipp11* locus |
| VPR_screening_R1 | CACAACCTGGTCAGTCCCTC | Primer for screening PYL1-VPR and ABI-LacI insertions at *Hipp11* locus |
| Fbn2_qPCR_F1 | AAATACTCCGAGGGCCCAAC | qPCR primer for mature *Fbn2* transcripts |
| Fbn2_qPCR_R1 | GGCGGGAACAGAATCCATCT | qPCR primer for mature *Fbn2* transcripts |
| Fbn2_qPCR_F2 | CGGAACAGCTTGTGTAGACCT | qPCR primer for mature *Fbn2* transcripts |
| Fbn2_qPCR_R2 | TGAAACCGGGCGGACATTTA | qPCR primer for mature *Fbn2* transcripts |
